# Expanding the toolbox of probiotic *Escherichia coli* Nissle 1917 for synthetic biology

**DOI:** 10.1101/2023.06.05.543671

**Authors:** Fang Ba, Yufei Zhang, Xiangyang Ji, Wan-Qiu Liu, Shengjie Ling, Jian Li

**Affiliations:** School of Physical Science and Technology, ShanghaiTech University, Shanghai, 201210, China

**Author notes:** Corresponding author. (J.L.) **Corresponding Author** Jian Li - School of Physical Science and Technology, ShanghaiTech University, Shanghai 201210, China.

**Keywords:** synthetic biology, probiotic engineering, *Escherichia coli* Nissle 1917, cryptic plasmid, integrase, cell-free protein synthesis

## Abstract

*Escherichia coli* Nissle 1917 (EcN) is a probiotic microbe that has the potential to be developed as a promising chassis for synthetic biology applications. However, the molecular tools and techniques for utilizing EcN have not been fully explored. To address this opportunity, we systematically expanded the EcN-based toolbox, enabling EcN as a powerful platform for more applications. First, two EcN cryptic plasmids and other compatible plasmids were genetically engineered to enrich the manipulable plasmid toolbox for multiple gene coexpression. Next, we developed two EcN-based enabling technologies, including the conjugation strategy for DNA transfer, and quantification of protein expression capability. Finally, we expanded the EcN-based applications by developing EcN native integrase-mediated genetic engineering capabilities and establishing an *in vitro* cell-free protein synthesis (CFPS) system. Overall, this study expanded the toolbox for manipulating EcN as a commonly used probiotic chassis, providing several simplified, dependable, and predictable strategies for researchers working in synthetic biology fields.

**For Table of Contents Use Only:** 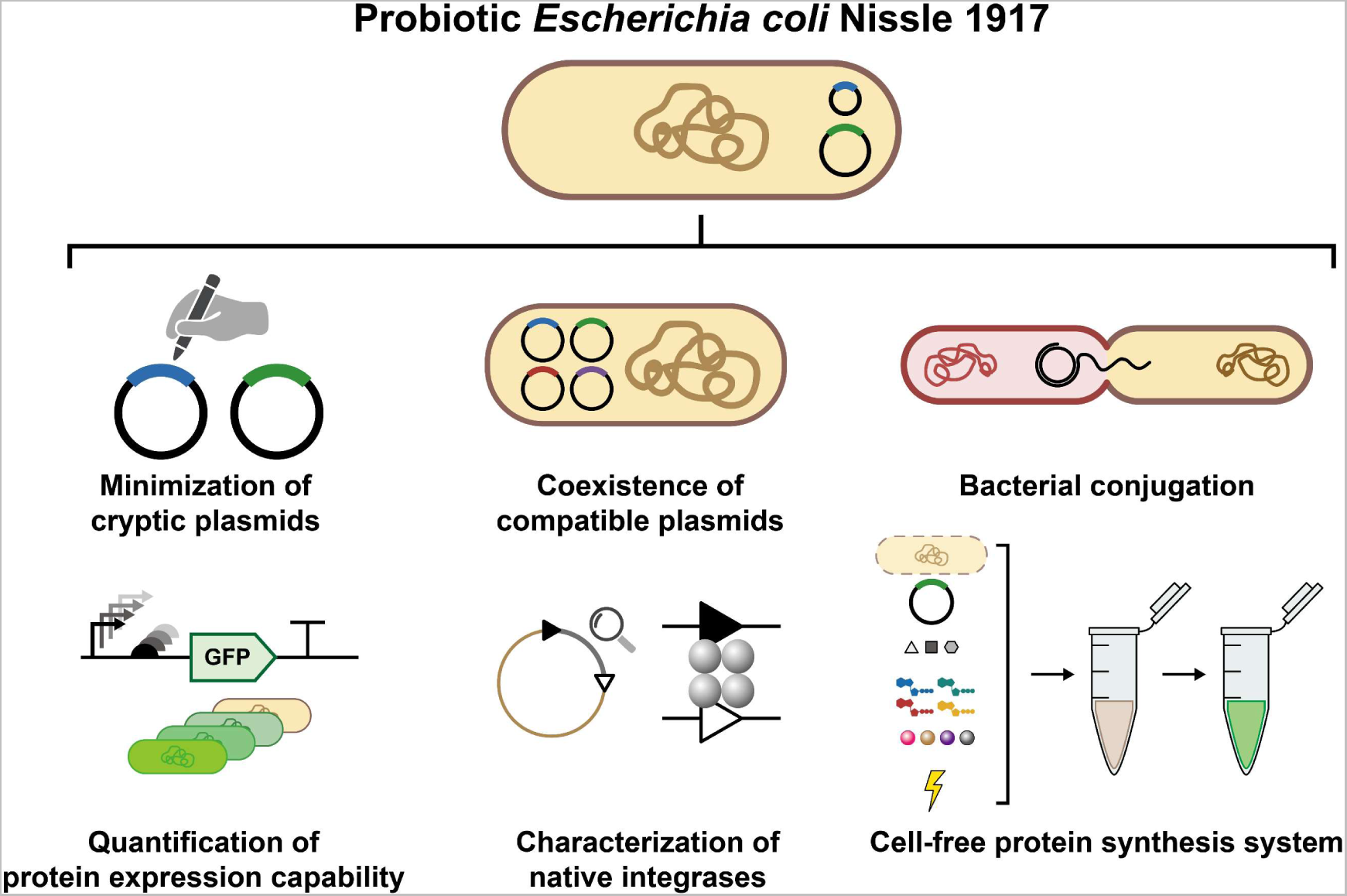

## Introduction

Microbes are widely used for molecule biosynthesis and biomanufacturing purposes.^1, 2^ Specifically, genetically engineered microbes can be used for the “Design-Build-Test-Learn” cycle,^3, 4^ including laboratory research,^5, 6^ industrial production,^7^ and biomedical living therapeutics.^8−10^ As a result, probiotics are excellent microorganism candidates due to their high-biosecurity, non-toxicity, biocompatibility, and well-acceptability.^10, 11^ Several probiotics, including gram-positive bacteria (e.g., *Bifidobacterium bifidum*,^12^ *Lactobacillus lactis*,^10, 13^ and *Streptococcus thermophilus*^14^) and gram-negative bacteria (e.g., *Bacteroides fragilis*^10^), have been isolated, engineered, and used as living factories. In addition, these chasses have been used in the biosensing,^5^ biomolecule production and delivery,^15^ metabolic regulation,^16^ antitumor therapy,^17^ and immunotherapy.^18^

Nonetheless, *Escherichia coli* Nissle 1917 (EcN) has been among the most widely used and acceptable strains since it was identified in 1917.^19^ Due to its unique superiority, such as its well-understood *Escherichia coli* background, long-term biosecurity record of human administration, great *in vivo* colonization ability, antagonistic, anti-inflammatory, and anti-invasive activity,^20^ EcN was utilized as both a laboratory chassis cell and a commercial probiotic product named “Mutaflor”.^19^ Up to now, EcN has become one of the most popular probiotic chassis for developing living therapeutics in pathogenic infections,^21^ metabolic regulation,^16^ GI tract inflammations,^15^ and antitumor treatments.^22^

Nowadays, some EcN-based genetic engineering strategies, including genome engineering (e.g., Lambda-Red recombination,^23^ integrase-based site-specific recombination,^24^ and CRISPR-Cas-based gene editing technologies^25−27^), cryptic plasmid engineering,^25−27^ and chassis modification,^24, 28^ have been designed, built, and tested. However, the genetic engineering toolbox of EcN can be further expanded in other fields due to its non-type strain properties.

To further meet these demands, EcN-based genetic engineering toolbox was systematically enriched to expand probiotic-based synthetic capabilities. First, EcN cryptic plasmids and other synthetic plasmids were modified to enrich the vector toolbox. In details, the two EcN cryptic plasmids pMUT1 and pMUT2 were successfully minimized, and the minimized pMUT2 origin of replication (oriV) was characterized as a 26 bp DNA sequence. Next, the coexistence and curing of up to four compatible plasmids were demonstrated, including oriVs of ColE1, ColE2, p15A, pSC101, and incW. Second, two EcN-enabling technologies were developed for DNA and protein manipulation. A simplified plasmid transfer strategy was developed (at least 30 minutes) via bacterial conjugation, which providing another efficient way than chemical or electrical transformation. Besides, EcN protein expression capability was quantified using three generalized genetic circuits, sfGFP expression ranged from 0.24 to 44.7 µg / mL. Third, two EcN-inspired applications were developed for potential probiotic functionalization. On the one hand, four new integrases in the EcN genome were first characterized, and the integrases were expressed and used for deletion, inversion, and integration depending on the designed *attP* / *attB* orientations. On the other hand, probiotic EcN-based cell-free protein synthesis (CFPS) system was successfully established (sfGFP yield reached approximately 160.8 µg / mL). In addition, lyophilized EcN CFPS systems retained acceptable activity within seven days.

We expanded the EcN-based genetic engineering toolbox for desired applications. It is worth emphasizing that four probiotic-derived integrases were first characterized as reliable genetic engineering tools for probiotic chassis modification. In addition, a probiotic-based CFPS system was successfully established for on-demand biomanufacturing of pharmaceutical proteins, vaccines, and drug production. Therefore, this study may inspire the engineering of more probiotics for broad applications in synthetic biology, metabolic engineering, and biomedical engineering.

## Results and Discussion

### Minimization of EcN Cryptic Plasmids pMUT1 and pMUT2

EcN contains two cryptic plasmids pMUT1 (3173 bp) and pMUT2 (5514 bp), which are not transmissible and without antibiotic resistance genes (**Figures S1A, B**).^25^ Although these plasmids have been modified and functionalized as synthetic vectors for specific purposes,^25−27^ they have not been comprehensively explored, such as their minimized size (**Figure 1A**). To address these issues, we initially analyzed plasmid sequences and annotations of putative open reading frames (ORFs) (**Figure S1B**). After deleting redundant sequences, pMUT1 was partially truncated from 3173 bp to 1312 bp (**Figure 1B**), while pMUT2 was truncated from 5514 bp to 3093 bp (**Figure 1C**). Next, the two truncated vectors were reconstructed with antibiotic resistance genes, and their existence stability was verified. Referring to previous reports, pMUT1 contains ColE1-like oriV, which is commonly derived as plasmid vectors (e.g., pET-series, pSB1C3, pJL1),^25^ but pMUT2 is mostly homologous to plasmid pUB6060 from *Plesiomonas shigelloides* and annotated as a ColE2-like oriV, which has not been comprehensively explored before.^25, 29^ Hence, to further minimized pMUT2 and precisely identified its oriV sequence, we performed truncation experiments and ensured its oriV sequence was 26 bp (**Figure S1C**). Lastly, to conveniently integrate gene of interests (GOI) into the truncated pMUT plasmids, we added BioBrick prefix and BioBrick suffix sequences into variants and finally constructed the minimized pMUT plasmids: pMUT1-mini (2423 bp) (**Figure 1B**) and pMUT2-mini (3136 bp) (**Figure 1C**).^30^ Furthermore, the modified plasmids were analyzed via agarose gel electrophoresis and correctly sequenced (**Figure 1D**, and **Figure S1D**). In summary, we successfully minimized two EcN cryptic plasmids, precisely identified ColE2-like oriV sequence, and constructed two pMUT-based minimized plasmids.

**Figure 1.**
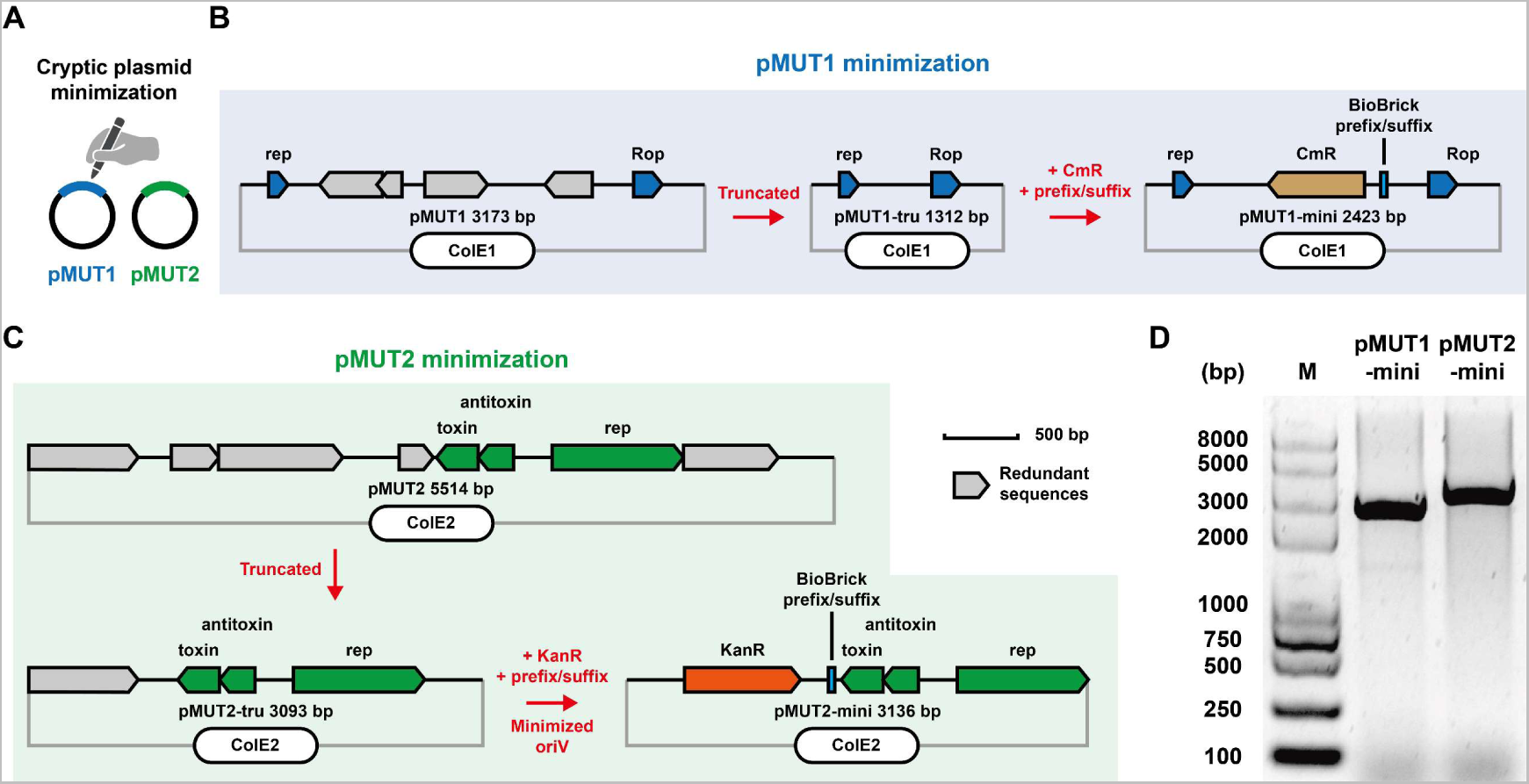
Minimization of cryptic plasmids. (A) Wild type EcN contains two cryptic plasmids pMUT1 and pMUT2. (B) Workflow of pMUT1 minimization. pMUT1-tru was truncated from pMUT1 by redundant sequence deletion, pMUT1-mini was derived from pMUT1-tru by adding antibiotic resistance gene CmR and BioBrick prefix / suffix sequences. (C) Workflow of pMUT2 minimization. pMUT2-tru was truncated from pMUT2 by redundant sequence deletion, pMUT2-mini was derived from pMUT2-tru by adding antibiotic resistance gene KanR and BioBrick prefix / suffix sequences. (D) Agarose gel electrophoresis of two minimized plasmids that linearized by FastDigest XbaI digestion.

### Coexistence and Curing of Compatible Plasmids in EcN

The maximum number of coexistent pairs of engineered plasmids was explored after the characterization of the compatible cryptic plasmids, notably, the oriV is crucial for plasmid existence in microbes. oriV is classified as different incompatibility groups depending on the replication mechanism, indicating that two different plasmids with the same oriV cannot coexist in one microbe because of the competence of the same replication machine since they may create an unpredictable environment.^31^ Therefore, establishing compatible plasmid vectors can expand the exogenous DNA capacity in one strain for synchronous gene expression. To meet these demands, a series of empty vectors (followed as iGEM BioBrick assembly standard^30^) mainly consisting of four parts (antibiotic resistance gene, oriV, BioBrick prefix / BioBrick suffix (for molecular cloning), and GOI) were developed (**Figure 2A**). The orthogonal and compatible plasmid groups in EcN were then assembled to follow the basic design that six plasmid vectors were constructed using four antibiotic resistance genes and five oriVs (**Figure 2B**, and **Figure S2**). When plasmids were successfully constructed in *E. coli* Mach1-T1, they were then transformed into wild type EcN and their copy number was quantified via qPCR. The results showed that pMUT2-mini (ColE2) had the highest copy number of ∼22 per cell, while pFB2 (pSC101) had the lowest copy number (∼2 per cell) (**Figure 2C**). Next, we gained four EcN strains that contain four compatible vectors, respectively. After seven days’ continuous culture, the EcNs did not lose the vectors and could normally grow on LB-agar plates (with four antibiotics) (**Figure 2D**). In addition, these vectors could be cured using spCas9-based gene editing tools as previously reported,^32^ which expanding the plasmid manipulation method in EcN. In conclusion, we expanded the compatible plasmid vectors for EcN and demonstrated, their coexistence capability for at least one week.

**Figure 2.**
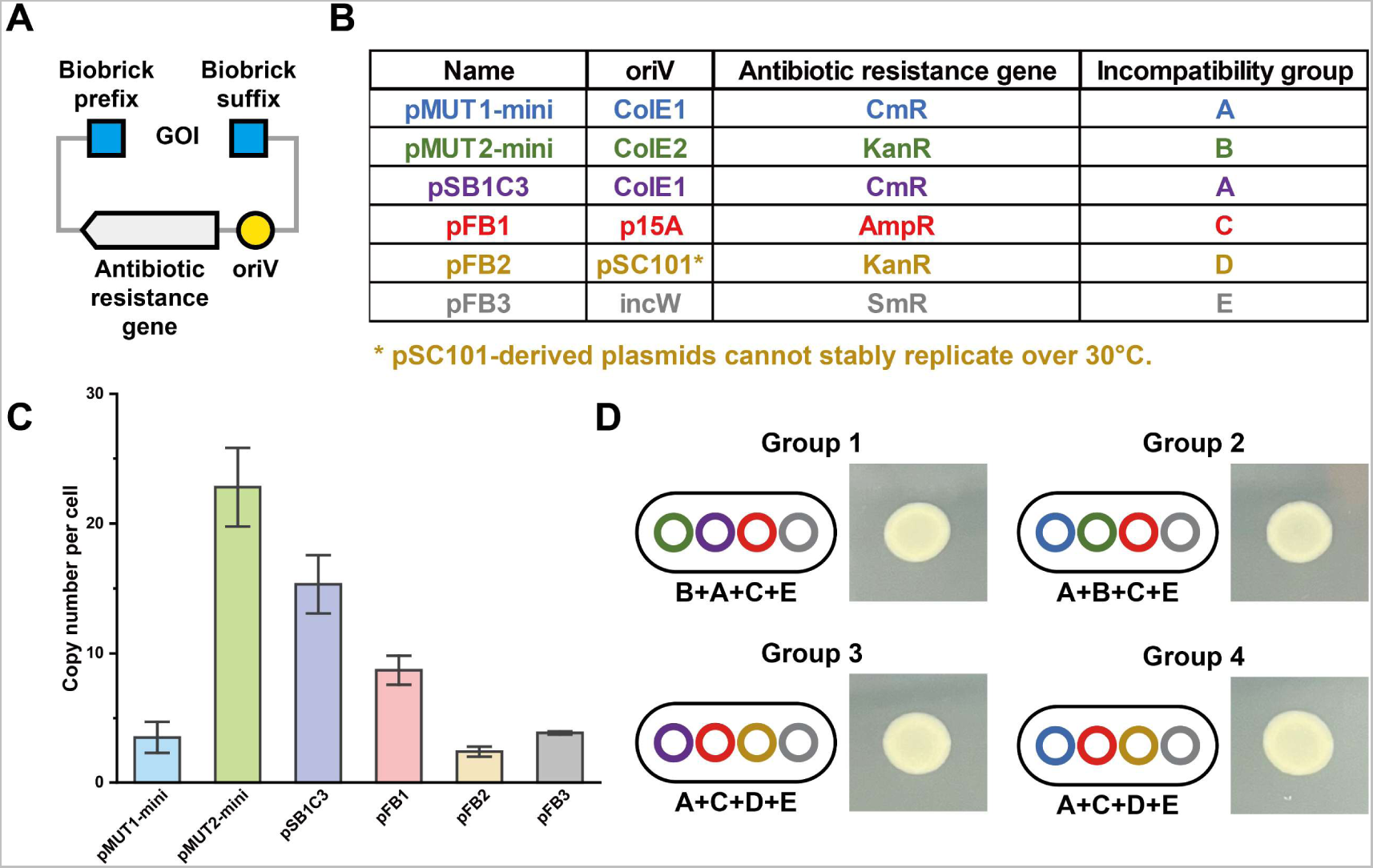
Coexistence of compatible plasmids in EcN. (A) Schematic design of compatible plasmid vectors. GOI (gene of interests) can be inserted into vectors by molecular cloning via BioBrick prefix / suffix sites. (B) Classification of vectors. (C) Copy number per EcN cell of six vectors. Each value (mean ± standard deviation, SD) was calculated with three biological replicates. (D) Compatible vectors were coexistent in EcN. All groups contained up to four compatible vectors in one strain, and could grow normally on LB-agar plates (with four antibiotics) after seven days’ cultivation. The experiments were performed as thrice with the same results.

### Bacterial Conjugation

So far, EcN-based chassis modification was performed by plasmid engineering. Next, we focused on the enabling technology to render EcN new properties and functionalities. It was worth noting that the EcN-based conjugation method for DNA transfer has not been commonly developed. Herein, a simplified strategy was developed for expanding EcN-based enabling technology.

Bacterial conjugation is one of the powerful tools of horizontal gene transfer within microbial communities, which enables donor strains to transfer their DNA through conjugative pili and the DNA was received by recipients.^33^ This strategy has been developed for bacteria-bacteria DNA transfer within different species.^34^ Depending on our previous attempts, the calcium chloride-based heat-shock transformation strategy was unsuccessful for EcN (data not shown). Besides, the efficiency of electroporation-based transformation is not high enough compared with the generally used laboratory *E. coli* strains, such as DH5α (**Figure S3**). Therefore, a simple, repeatable, and widely accepted plasmid transfer method for EcN is necessary. Herein, *E. coli* S17-1 λpir (ATCC BAA-2428) was selected as the donor cell because its conjugative pili system (RP4-2) is integrated into the genome for RP4-oriT-based DNA conjugation transfer.^35, 36^ EcN was used as a recipient (harboring control plasmid to provide another antibiotic resistance gene for transconjugant selection). After performing the conjugation workflow (**Figure 3A**), two donor plasmids (compatible with EcN control plasmids) were successfully transferred into recipient EcN strains, with conjugation efficiency of up to 10^5^ CFU per 1 mL OD_600_ = 1 recipient cell (within 30 min) (**Figures 3B, C**). However, conjugation efficiency did not significantly increase after 30 min, indicating that EcN-based conjugation is a powerful and time-saving DNA transfer strategy.

**Figure 3.**
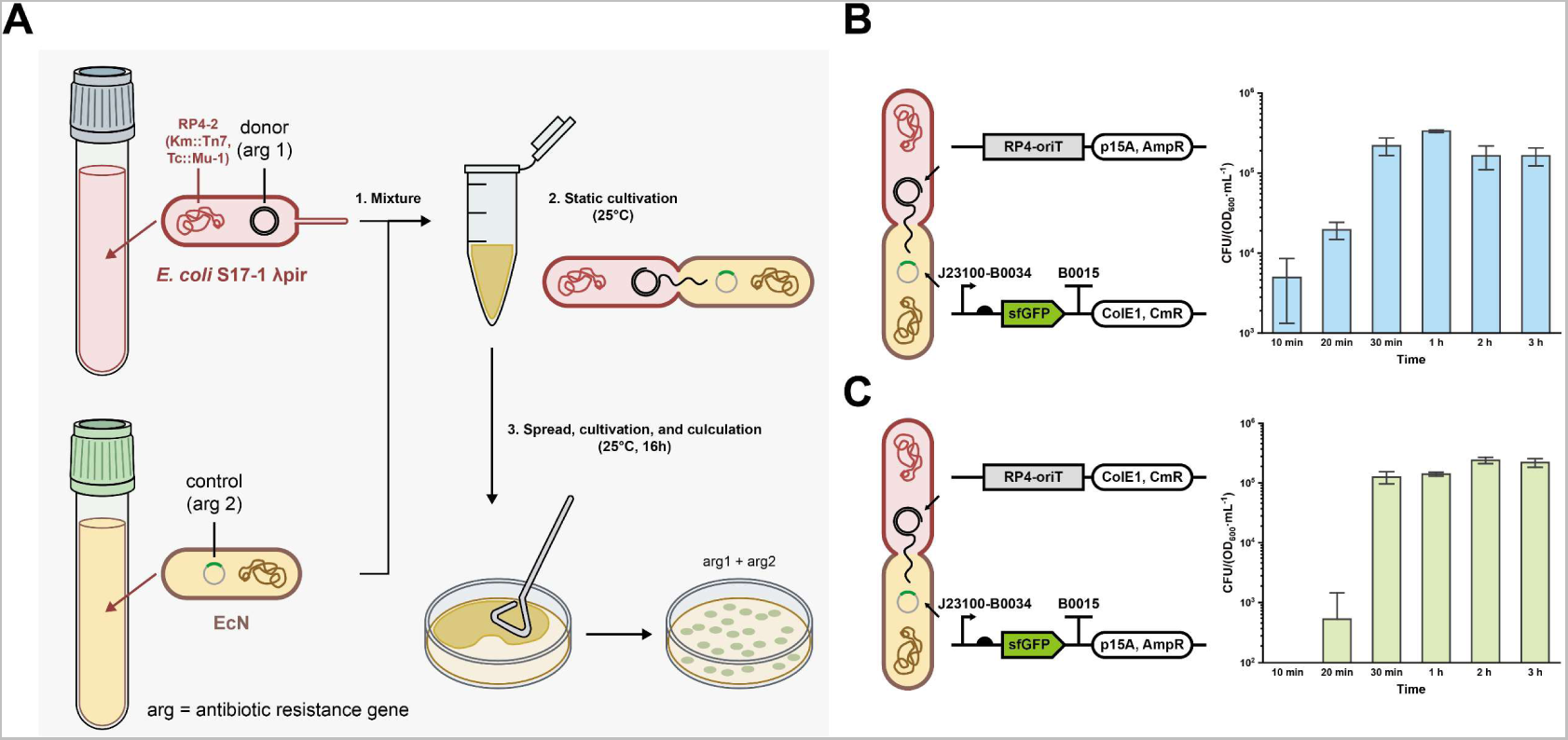
Bacterial conjugation. (A) Workflow of “*E. coli* S17-1 λpir - EcN” conjugation. (B) Conjugation efficiency of plasmid pFB345 (pFB1-RP4-oriT) transferred into EcN (harboring pFB346 as control plasmid). (C) Conjugation efficiency of plasmid pFB347 (pSB1C3-RP4-oriT) transferred into EcN (harboring pFB348 as control plasmid).

### Protein Expression Capability of EcN

Probiotic EcN has been widely used for protein expression for therapeutic applications, such as bacterial infections,^21^ inflammatory bowel disease,^18^ metabolic diseases,^16^ and antitumor immunotherapy.^22^ A higher protein expression capability is necessary because proteins act as therapeutic drugs or catalytic enzymes for metabolite production. Hence, we next attempt to explore the protein expression capability of EcN, here we select the superfolder green fluorescent protein (sfGFP) as the reporter. First, the protein expression capability of four *E. coli* strains with five constitutive σ70 promoters was compared (**Figure 4A**),^37^ however, EcN had the lowest sfGFP expression level, with the highest yield of ∼ 17 µg per 1 mL OD_600_ = 1 culture (**Figure 4A, right**). Then, an arabinose-inducible sfGFP expression plasmid (with different arabinose concentrations) was then established, obtaining the highest sfGFP yield of ∼ 8 µg per 1 mL OD_600_ = 1 culture (**Figure 4B**). Furthermore, the strong T7 RNA polymerase-based protein expression systems with two compatible plasmids (one supplying low constitutive T7 RNA polymerase and the other one was constructed using T7 promoter and lacI-lacO transcriptional repressor system responding to isopropyl β-D-1-thiogalactopyranoside (IPTG) induction) were used to improve the expression capability (**Figure 4C**).^24, 38^ These two kinds of plasmids were coexistent in EcN, when IPTG was not added in LB medium, the leaked T7 RNA polymerase caused a considerable sfGFP yield (J23113-B0034, ∼ 41 µg per 1 mL OD_600_ = 1).^38^ When 0.5 mM was added, the highest induced sfGFP expression reached ∼ 44 µg per 1 mL OD_600_ = 1. In summary, three sfGFP expression systems showed that the range of EcN protein expression capability is from 0.24 to 44.7 µg per 1 mL OD_600_ = 1 (**Figure 4D**), providing references for EcN-based protein expression with biomanufacturing applications.

**Figure 4.**
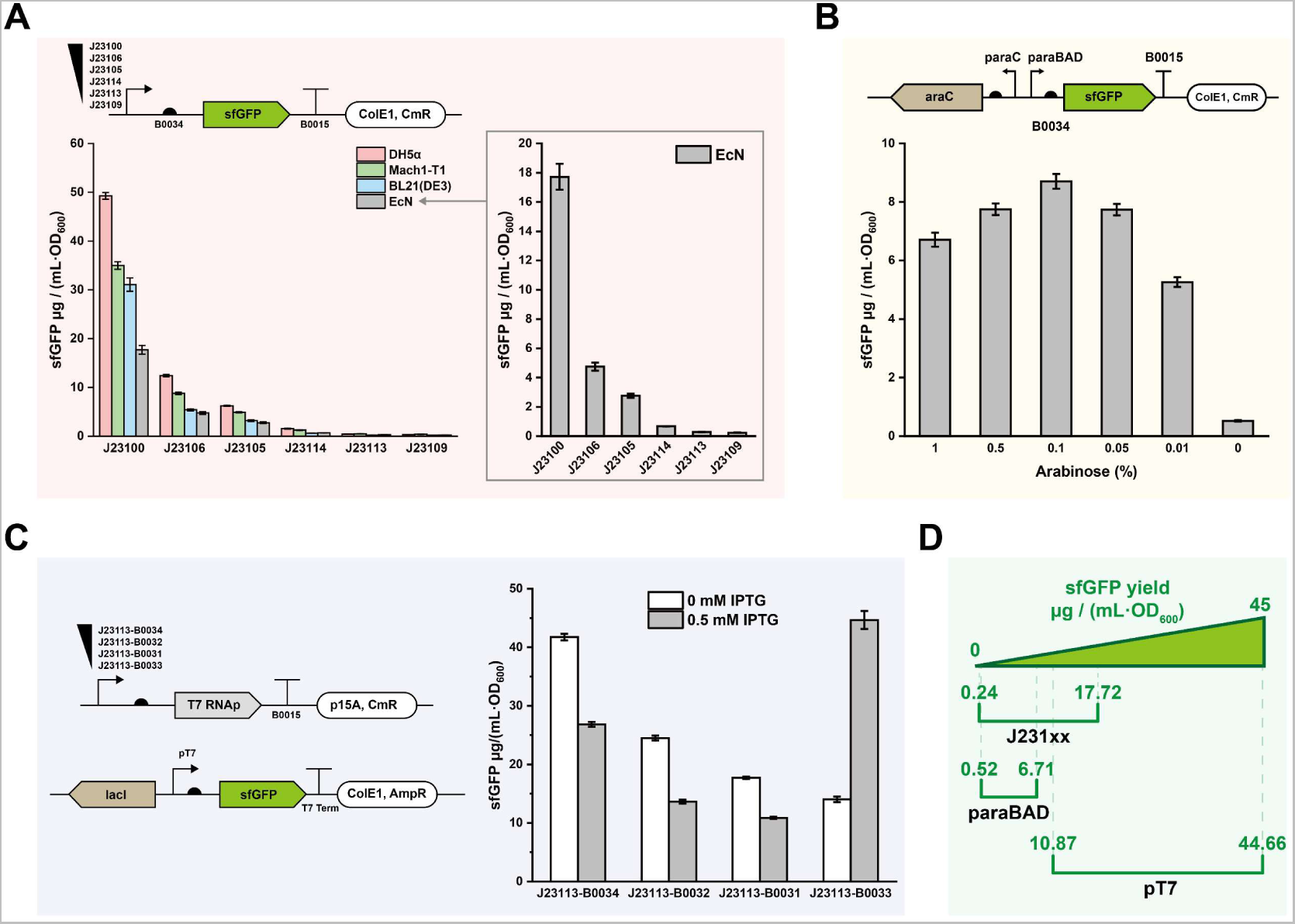
Quantification of EcN protein expression capability. (A) Four *E. coli* strains (K-12 strain DH5α, W strain Mach1-T1, B strain BL21(DE3), and EcN) and six sfGFP expression plasmids (pFB346, pFB349-pFB353) were used. (B) Arabinose-induced sfGFP expression plasmid pFB354 was used in EcN. (C) T7 RNA polymerase expression plasmids (pFB355-pFB358) and IPTG-induced sfGFP expression plasmid (pFB359) were coexisted in EcN. (D) sfGFP yield of all three EcN sfGFP expression systems. Each value (mean ± standard deviation, SD) was calculated with three biological replicates.

### Characterization and Functionalization of EcN Native Integrases

Up to now, we performed the chassis modification and enabling technology expansion of EcN, therefore, it could be further explored as powerful platform for both *in vivo* and *in vivo* applications. Depending on our previous studies, the discovery of native integrases from EcN caught our eyes because they might be widely expanded the genetic engineering applications for defined useful purposes.^39^

Integrases are a class of enzymes that catalyze DNA-DNA site specific recombination between bacteriophage and host microbes. Bacteriophage DNA can be integrated into the host cell genome as prophage.^40^ Recently, integrases have been widely developed as genetic tools in synthetic biology fields.^41^ However, most of the reported integrases were isolated from non-probiotic microbes that their biosafety and acceptability were limited. Fortunately, our preliminary investigation suggested nine potential integrases in EcN genome (**Figure S4**). After analysis of comparative genome annotation (**Figure S5**), we discovered four potential prophage sequences (each prophage contains one integrase) (**Figures 5A**, **B**). Next, four arabinose-induced integrase expression plasmids were constructed to test the *in vivo* expression ability (**Figure 5C**). Notably, InterPro analysis suggested the four integrases’ classification as tyrosine integrase family (**Figure S5**).^42^ As a result, 6×HisTag was added in the N-terminal (DNA-binding domain) rather than C-terminal (catalytic domain) to ensure the catalytic activity. Like the other previously reported integrases, these four EcN native integrases could be expressed as acceptable solubility (**Figure 5D**, and **Figure S6**) and purified from *E. coli* cells (**Figure 5E**). Afterwards, the *attP* and *attB* sites were then reassembled (**Figure 5F**, and **Figure S7**), and plasmid was constructed to characterize the integrase activity (**Figure 5G**). As expected, four integrases were able to catalyze recombination events (**Figure 5H**). Furthermore, to ensure the C-terminal catalytic residues, C-terminal tyrosine residues were converted into alanine or phenylalanine.^43^ As the InterPro annotated, the key tyrosine residues (int 1 395Y, int 2 384Y, int 3 379Y, and int 4 374Y) are essential for recombination (**Figure S8**). To sum up, we identified four EcN native integrases, successfully characterized their recombination activity, and acquired reassembled attachment sites.

**Figure 5.**
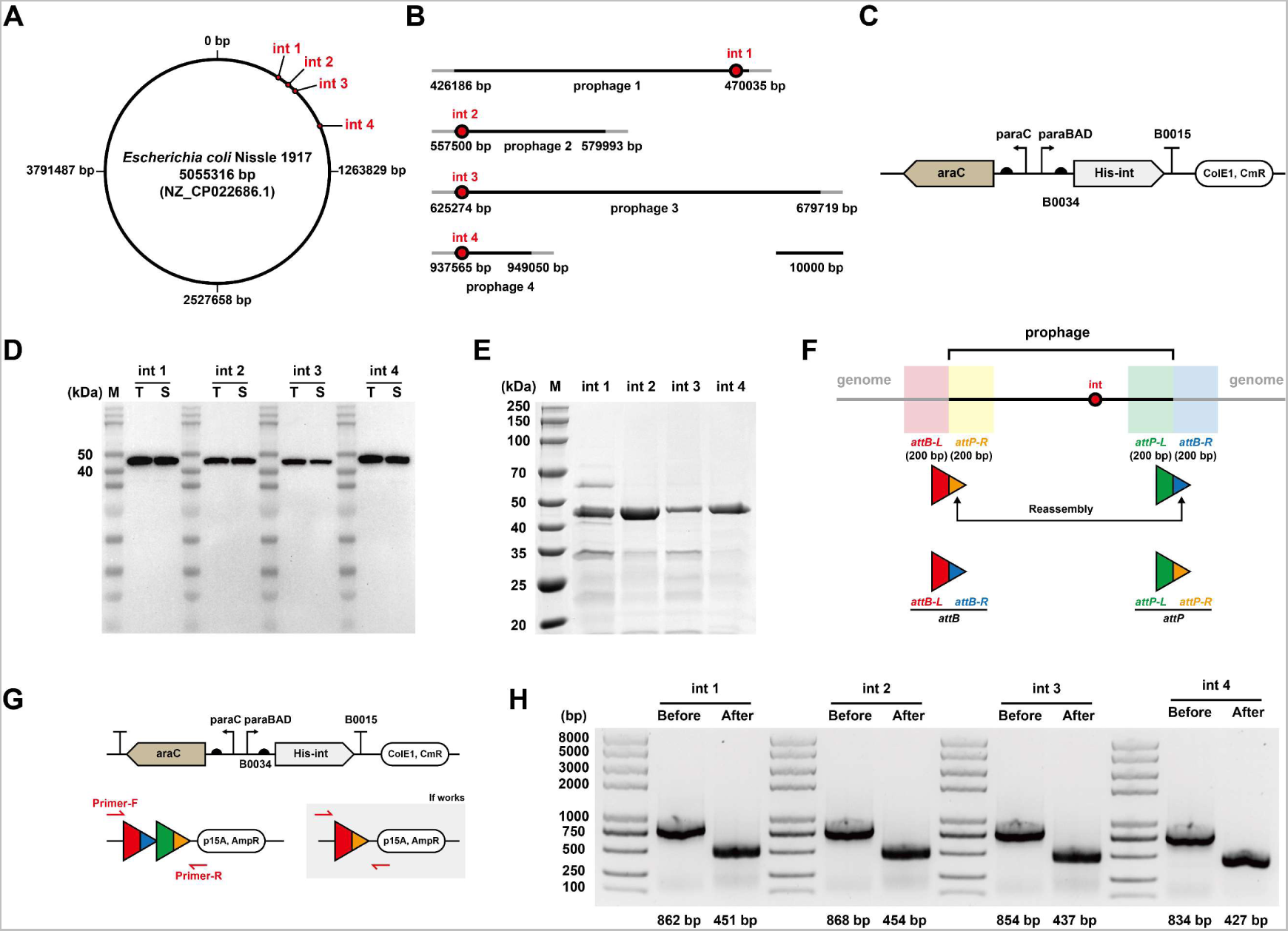
Characterization of EcN native integrases. (A) Four potential integrases and their localization in EcN genome. (B) Detailed prophage and integrase locations in EcN genome. (C) Schematic of four arabinose-induced integrase expression plasmids (pFB360-pFB363). (D) Western blot analysis of four integrases with N-terminal 6×HisTag (int 1: 50.2 kDa, int 2: 46.7 kDa, int 3: 49.6 kDa, int 4: 47.6 kDa). M: marker, T: total, S: soluble. (E) SDS-PAGE analysis of four purified integrases. (F) Workflow of integrase *attB* / *attP* reassembly strategy. (G) Schematic design of integrase activity test. int 1 as an example: *E. coli* Mach1-T1 harboring both int 1 expression plasmid pFB360 and *attP* / *attB* sites plasmid pFB368 was cultured, PCR product size indicated whether int 1 performed activity. (H) Characterization results of all four integrases. Before: PCR product without integrase catalysis, After: PCR product with integrase catalysis. All the experiments were performed as thrice with the same results.

DNA fragments can be deleted, inverted, and integrated by integrases depending on the orientation of *attP* / *attB.* Herein, the four integrases were functionalized with different *attP* / *attB* orientations. First, the reporter plasmids were designed as constitutive sfGFP expression within the same direction as *attP* / *attB* pairs to assess deletion efficiency, *E. coli* will lose fluorescence if the sfGFP is deleted by integrase (**Figure 6A**). Results showed that the four integrases had 100% deletion efficiency. Second, the reporter plasmids were designed as constitutive sfGFP expression within the opposite direction of *attP* / *attB* pairs to evaluate the inversion efficiency. A desired PCR product can be obtained if DNA is inverted, and *E. coli* may also not lose fluorescence (**Figure 6B**). Agarose gel electrophoresis results showed that the four integrases had 100% inversion efficiency (**Figure S9**). Third, to perform the integrase-mediated DNA integration, at the beginning, all four *attB* sites were initially integrated into *E. coli* MG1655 genome, then the donor plasmids were constructed with *attP* site, antibiotic resistance gene (SmR), and R6K-oriV (cannot replicate without gene *pir*)^44^ (**Figure S10**). The experiments were performed with a standard protocol (**Figure 6C**), and the integration, efficiency was measured from 10^3^ to 10^1^ (**Figure 6D, right**). As expected, all four EcN-derived integrases were successfully functionalized as three performances: deletion, inversion, and integration.

**Figure 6.**
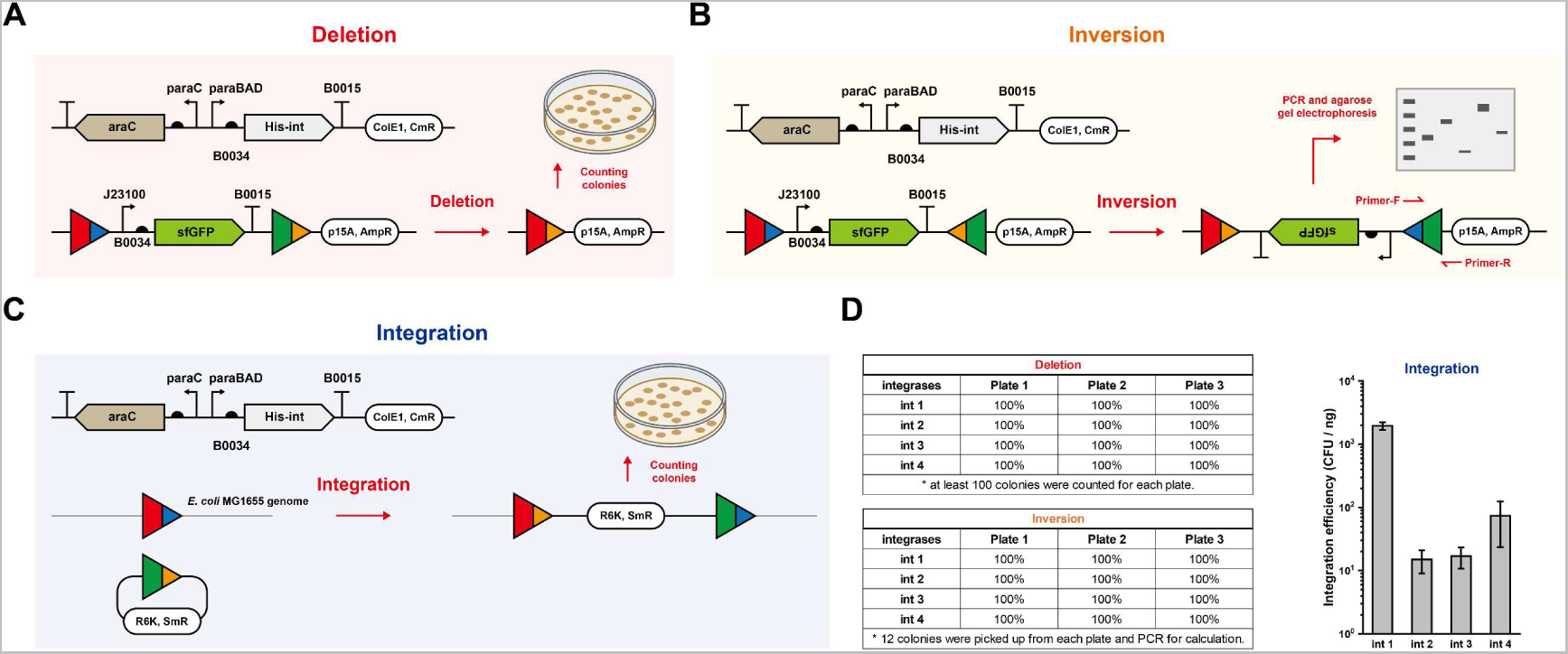
Functionalization of EcN native integrases. Depending on the orientation of *attP* / *attB* sites, integrases worked as (A) deletion, (B) inversion, or (C) integration. (D) Efficiencies of deletion, inversion, and integration. Each value (mean ± standard deviation, SD) of integration was calculated with three biological replicates.

### Establishment of the Probiotic EcN-Based CFPS System

CFPS systems have been widely developed for *in vitro* protein synthesis with the advantages of time-saving, easily manipulable, microscale, high-throughput, and portable.^45^ CFPS has also inspired the multidiscipline research fields, including genetic engineering,^46^ metabolic engineering,^47−50^ high-throughput screening,^51^ pharmaceutical production,^52, 53^ and education.^54, 55^ Therefore, a powerful chassis is essential for CFPS system establishment. Several new microbial chasses, including *Pichia pastoris*,^56^ *Bacillus subtilis*,^57^ *Streptomyces*,^46, 58−60^ *Pseudomonas putida*,^61^ *Vibrio natriegens*,^62, 63^ and *Klebsiella pneumoniae*,^64^ have been recently developed as CFPS systems. However, probiotic-based CFPS systems, which have great biosafety and higher acceptability, have not been widely developed. Thus, we aimed to develop a probiotic EcN-based CFPS system to expand chassis selection (**Figure 7A**). Initially, we supposed to simplify the EcN genetic engineering strategy to avoid complicated genetic manipulation. To meet the high-level transcription demands, we constructed a series of plasmids for T7 RNA polymerase constitutive expression rather than integrating T7 RNA polymerase into EcN genome (**Figure 7B**). SDS-PAGE suggested the soluble expression of T7 RNA polymerase *in vivo* (**Figure 7C**). Next, EcN crude extract for CFPS reactions was prepared based on the *E. coli* crude extract preparation procedure with optimized parameters.^65, 66^ The highest sfGFP yield was 160.8 ± 6.6 µg / mL with J23114 promoter at 30 °C (**Figures 7D, E**). Furthermore, the lyophilized CFPS mixture was stored at room temperature for days (**Figure 7F**), and rehydrated EcN CFPS system could produce considerable sfGFP within one week (**Figure 7G**), which providing a portable probiotic-based biomanufacturing platform for various applications.

**Figure 7.**
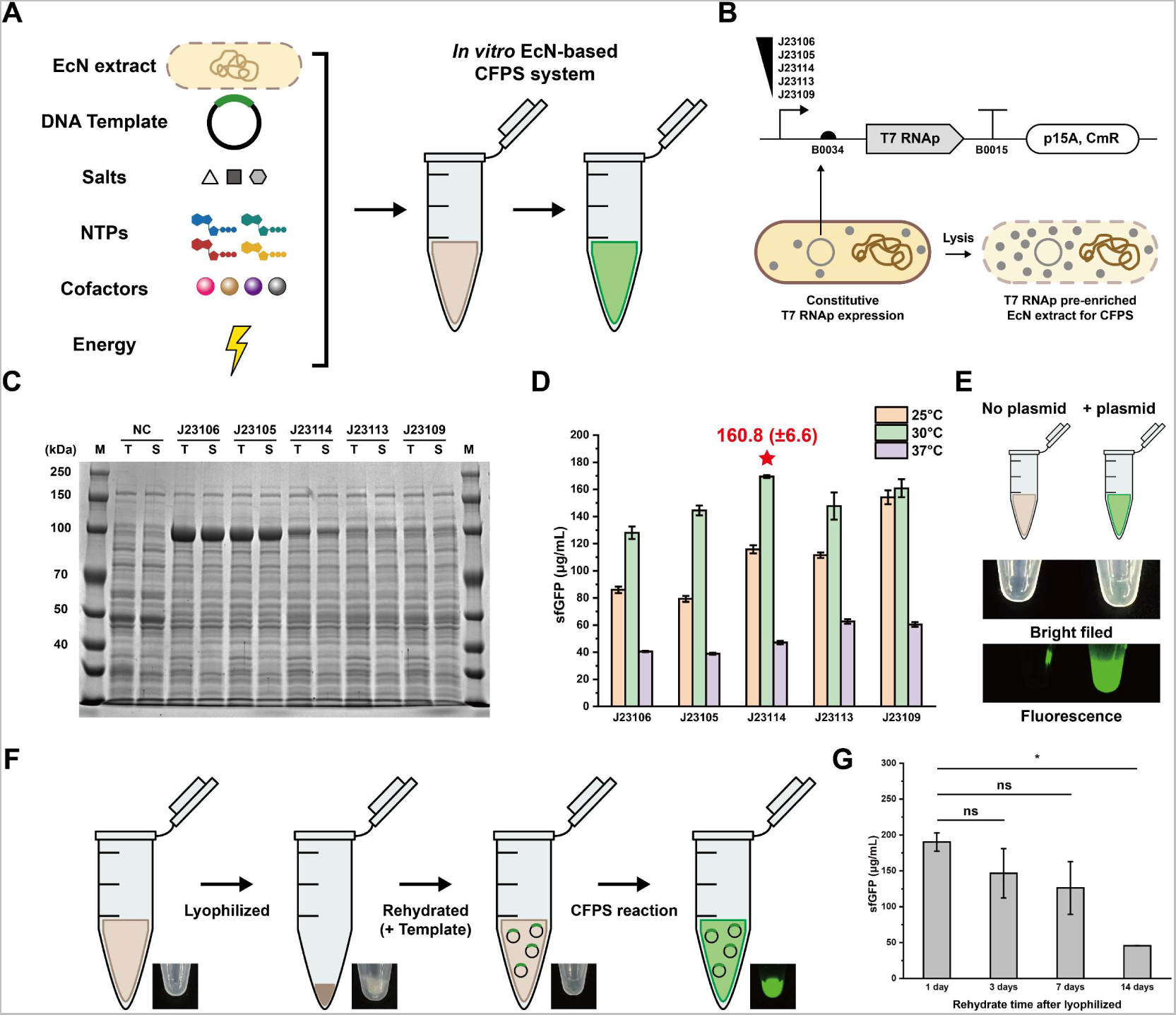
EcN-based cell-free protein synthesis (CFPS) system. (A) Schematic diagram of EcN-based CFPS system. (B) EcN strains (harboring T7 RNA polymerase constitutive expression plasmids pFB384- pFB388) for preparing CFPS crude extracts. (C) SDS-PAGE analysis of EcN strains in (B), T7 RNA polymerase (98.9 kDa) were clearly observed. M: marker, T: total, S: soluble. (D) sfGFP expression yield of five EcN strains and under three different temperatures for 3 h. (E) Visualization of one 15 µL standard CFPS reaction (J23114-T7 RNA polymerase, 30 °C) under bright field and fluorescence imager. (F) Workflow of EcN CFPS system lyophilization and rehydration. (G) sfGFP expression yield of lyophilized EcN CFPS reactions after 1 day, 3 days, 7 days, and 14 days. Each value (mean ± standard deviation, SD) of integration was calculated with three biological replicates. Student’s *t*-tests were used for statistical analysis, and *P* < 0.05 indicates statistical significance (* *P* < 0.05, “ns” means not significant).

## Conclusions

As synthetic biology quickly emerged in the last twenty years, probiotics have been gradually focused as one kind of desired chassis for useful purposes.^10^ Meanwhile, expanding the genetic toolbox is necessary for engineering more non-model probiotics that rarely explored.^6^ In this study, EcN-based toolbox was successfully expanded from three aspects. First, we focused on the chassis modification in two ways: two EcN-based cryptic plasmids were minimized and modified as BioBrick vectors, and plasmid coexistence in EcN was extended to four compatible groups. Second, enabling technologies were developed as two methods: simplified EcN-based conjugation strategy, and capability exploration of protein expression. Third, to enrich the EcN-based applications, four EcN native integrases were successfully identified, characterized, and functionalized into genetic circuit design and genome modification *in vivo*. Furthermore, EcN-based CFPS system was established as a probiotic-based *in vitro* biomanufacturing platform.

Overall, EcN toolbox was expanded through genetic modification, DNA transfer, protein expression capability quantification, new enzyme characterization, and CFPS system establishment, which may provide convincible strategies for future probiotic engineering and related studies. Notably, two cryptic plasmids were first minimized, followed by the identification of the ColE2-oriV sequence (26 bp in pMUT2). Four probiotic-derived integrases and their activity were also assessed, enriching the integrase toolbox for genetic circuit design and gene modification purposes. Besides, this probiotic-based CFPS system may expand the *in vitro* protein expression platform with higher biosecurity, non-toxicity, and well-acceptability.

Looking forward, we expect this study can inspire the engineering of more non-model or probiotic microbes to expand toolbox for broad applications in synthetic biology, genetic engineering, metabolic engineering, and biomedical engineering.

## Methods

### Strains, Vectors, Plasmids, crRNAs, Primers, and Reagents

The details of *E. coli* strains, vectors, plasmids, crRNAs, and primers used in this study are listed in **Tables S1** and **S3-S6**, **Supporting Information**. Some vectors and plasmids were derived from the author’s previous works.^39, 67^ The complete sequences of all plasmids are listed in **Supporting Information** (**Excel Sheet Data**), and their correctness was verified by Sanger sequencing (GENEWIZ) unless otherwise noted. Q5 High-Fidelity DNA polymerase (New England Biolabs), Phanta Super-Fidelity DNA polymerase (Vazyme), FastPure Gel DNA Extraction Mini Kit (Vazyme), and pEASY - Basic Seamless Cloning and Assembly Kit (TransGen Biotech) were used for molecular cloning. DreamTaq Green PCR Master Mix (Thermo Fisher Scientific) was used for colony PCR. FastDigest Restriction Enzymes (Thermo Fisher Scientific) were used for plasmid linearization. Lysogeny broth (LB) liquid medium contained 10 g tryptone, 5 g yeast extract, and 10 g sodium chloride in 1 L ddH_2_O. LB-agar plates were prepared by adding 15 g agar per liter LB liquid medium. Antibiotic stocks (1000×, dissolved in ddH_2_O unless otherwise noted) were 100 mg / mL ampicillin, 50 mg / mL kanamycin, 34 mg / mL chloramphenicol (dissolved in ethanol), and 50 mg / mL streptomycin.

### Genetic Parts

All genetic parts used in this study are listed in **Table S2**, **Supporting Information**. Genetic parts shorter than 150 bp were synthesized within the oligonucleotides (GENEWIZ) and inserted as DNA fragments during molecular cloning. Some genetic parts were derived from the author’s previous works.^39, 67^ EcN-based DNA sequences (including cryptic plasmids pMUT1 / pMUT2, four integrases, and integrases-associated *attP* / *attB* sites) were gained via PCR amplification referring to EcN genome sequence (GenBank: NZ_CP022686.1). RP4-oriT sequence was referred to GenBank: X14165.1. All genetic parts were checked for correctness with Sanger sequencing.

### Plasmid Construction

All plasmids were constructed by Gibson Assembly strategy. In brief, all linear DNA fragments were purified by gel extraction and then assembled by using pEASY - Basic Seamless Cloning and Assembly Kit. After assembly, the reaction mixture was added to 50 µL competent cells for transformation, which were incubated overnight on LB-agar plate (with antibiotics for selection). After colony PCR, probable products were sequenced by Sanger sequencing.

### Cryptic Plasmid Minimization

DNA sequences of pMUT1 (GenBank: CP058218) and pMUT2 (GeneBank: CP058219) were gained via PCR amplification (EcN whole-cell as PCR template). Minimization was performed referring to both gene annotations and previous reports.^25^ pMUT1-mini was constructed from three DNA fragments: pMUT1-tru (partial sequence of pMUT1), CmR gene (from pSB1C3), and BioBrick prefix / BioBrick suffix (oligonucleotides). pMUT2-mini was constructed from three DNA fragments: pMUT2-tru (partial sequence of pMUT2), KanR gene (from pKD4),^68^ and BioBrick prefix / BioBrick suffix (oligonucleotides). pMUT2-based 26 bp ColE2-oriV sequence was characterized by *in vivo* incremental truncation process. In brief, series of truncated (200 bp, 20 bp, 4 bp, 1 bp, in steps) plasmid variants were constructed and transformed into *E. coli* Mach1-T1 competent cells to determine the 5’ site and 3’ site of oriV. (non-survival plasmid variants lack complete oriV site for plasmid replication).

### Compatible Plasmid Coexistence, Plasmid Copy Number Calculation, and Curing in EcN

Each plasmid was transformed into EcN via electroporation, respectively. The EcN strains with four compatible plasmids were inoculated on LB-agar plates (with four antibiotics) and cultured overnight (37 °C, 16 h) to check the colony formation.

The copy number of each plasmid in EcN was calculated via qPCR,^39^ 16s rRNA genome sequence (six copies in one cell) was used as reference. The copy number per EcN cell was calculated as “6 × antibiotic resistance gene copy number / 16s rRNA copy number”.

Plasmid curing was performed as previously reported strategies,^32^ in brief, the pSC101 oriV-derived plasmid pFB2 (high temperature sensitive) was cured by over 30 °C continuous cultivation, ColE1 oriV-derived plasmids were cured by pCas, p15A and incW-oriV derived plasmids were cured by “pCas + crRNA plasmids (pFB390 - pFB395)”.

### Bacterial Conjugation

Donor strains *(E. coli* S17-1 λpir, harboring plasmid with RP4-oriT sequence) and recipient strains (EcN, harboring plasmid for antibiotic selection) were cultivated in 5 mL liquid LB medium (with antibiotics) at 37 °C and 250 rpm for 16 h. On the second day, for each culture, 1 mL mixture (adjusted OD_600_ to ∼1) was transferred into a 1.5 mL tube and centrifuged at 10000 g for 1 min. Then the pellets were resuspended by 1 mL LB medium (room temperature 25 °C, without any antibiotics) and centrifuged at 10000 g for 1 min, repeated the above wash step for another two times, and then resuspended the pellets by 1 mL LB medium (room temperature 25 °C, without any antibiotics). Afterwards, transferred both 50 µL donor strain culture and 50 µL recipient strain culture into a new 1.5 mL tube as total 100 µL, then steadily placed the mixture at 25 °C for hours. When conjugation finished, vortexed the mixture and added 900 µL LB medium (room temperature 25 °C, without antibiotics) for dilution. Finally, took 100 µL diluted mixture onto LB-agar plates (with two antibiotics) and cultured at 37 °C for 16 h. On the third day, counting the colonies (CFU) for conjugation efficiency calculation.

### *In vivo* Fluorescence Quantification

*E. coli* strains were cultivated in 5 mL liquid LB medium (with antibiotics, and with arabinose or IPTG if necessary) at 37 °C and 250 rpm for 16 h. On the second day, for each culture, 1 mL mixture (adjusted OD_600_ to ∼1) was transferred into a 1.5 mL tube and centrifuged at 10000 g for 1 min. Then the pellets were resuspended by 1 mL 1× PBS (Phosphate-buffered saline, pH 7.4), and sfGFP fluorescence was measured by microplate reader (Synergy H1, BioTek) and performed with excitation and emission wavelengths at 485 and 528 nm, respectively. Depending on the standard curve of “sfGFP (µg / mL) - Fluorescence (a.u.)” (**Figure S11**, **Supporting Information**), the *in vivo* sfGFP yield was normalized as “µg / (mL·OD_600_)”.

### Integrase Identification and Characterization

Potential EcN-based native integrases were identified via comparative genome annotations.^28^ After the initial round of integrase searching, nine candidates (**Figure S4**, **Supporting Information**) were identified and classified by InterPro.^42^ Next, the supposed attachment sites (both *attL* and *attR*) were analyzed, and gained relying on the potential prophage location and short DNA repeats (< 20 bp, DNA core sequences for integrases recombination). After reassembly *attL* / *attR* to *attP* / *attB*, plasmids were constructed for functional verification of “integrase - *attP* / *attB*” pairs.

### Expression and Purification of Integrases

The procedure was the same as our previous study.^39^ The difference is that pFB360- pFB363 were used in *E. coli* Mach1-T1 for integrase expression, and the inducer was replaced by 1% w/v arabinose than 0.5 mM IPTG.

### Tyrosine Residue Mutation of Integrases

Depending on the InterPro results (**Figure S5**, **Supporting Information**), the C-terminal tyrosine residues were replaced by alanine or phenylalanine.^43^ The tyrosine residue mutation plasmids were derived from pFB360-pFB363, mutants were characterized as same as wild type.

### Functionalization of Integrases

Here we describe integrase 1 as an example, the other three integrases were performed similarly.

Deletion: *E. coli* Mach1-T1 (harboring pFB364) was transformed with pFB360, on the second day, the colonies on LB-agar plates were picked up and inoculated into 5 mL liquid LB medium (with two antibiotics and 1% w/v arabinose) at 37 °C and 250 rpm for 16 h. On the third day, the culture was streaked onto LB-agar plates and cultured overnight at 37 °C for 16 h. On the fourth day, the colony’s fluorescence was observed under UVP ChemStudio (analytikjena), and the integrase 1-based deletion efficiency was calculated as “nonfluorescent colony number / total colony number”.

Inversion: *E. coli* Mach1-T1 (harboring pFB368) was transformed with pFB360, on the second day, the colonies on LB-agar plates were picked up and inoculated into 5 mL liquid LB medium (with two antibiotics and 1% w/v arabinose) at 37 °C and 250 rpm for 16 h. On the third day, the culture was streaked onto LB-agar plates and cultured overnight at 37 °C for 16 h. On the fourth day, the colonies were picked up and performed colony PCR (forward primer: int 1-inversion-F, reverse primer: VR, **Supporting Information**), if PCR product of the colony matched correct size (also this colony did not lose fluorescence), this colony was successfully performed the integrase 1-based inversion.

Integration: *E. coli* MG1655 (integrase 1 *attB* site has been inserted into the genome) harboring pFB360 was inoculated into 5 mL liquid LB medium (with 1% w/v arabinose) at 37 °C and 250 rpm for about 3 h, when OD_600_ reached 0.6, the cell was washed by ddH_2_O thrice and prepared as electroporation competent cell (concentrated 10 times, OD_600_ = ∼6). Then 1 ng donor plasmid pFB380 was electro-transformed into 50 µL competent cell, separated onto LB-agar plates (with two antibiotics), and cultured overnight at 37 °C for 16 h. On the second day, the colony number (CFU) was counted, and the integrase 1-based integration efficiency was calculated as CFU / ng (donor plasmid).

### Preparation of EcN Cell Extracts

Cell cultivation, harvest, and lysis were prepared according to our previous reports.^66^ Some parameters were specially optimized for EcN Cell Extracts, the revised workflow is shown in **Supporting Methods** (**Supporting Information)**. The final cell extracts were stored at −80 °C until use.

### Cell-Free Protein Synthesis (CFPS) and Protein Analysis

Standard CFPS reactions were performed in 1.5 mL tubes with a total volume of 15 µL. Each CFPS reaction contained: 12 mM magnesium glutamate, 10 mM ammonium glutamate, 130 mM potassium glutamate, 1.2 mM ATP, 0.85 mM each of GTP, UTP, and CTP, 34 µg / mL folinic acid, 170 µg / mL of *E. coli* tRNA mixture, 2 mM each of 20 standard amino acids, 0.33 mM nicotinamide adenine dinucleotide (NAD), 0.27 mM coenzyme A (CoA), 1.5 mM spermidine, 1 mM putrescine, 4 mM sodium oxalate, 33 mM phosphoenolpyruvate (PEP), 13.3 µg / mL plasmid, and 27% (v/v) of cell tract. CFPS reactions were incubated for at least 3 h, and at 30 °C (unless otherwise noted). Cell-free expressed sfGFP was measured by microplate reader (Synergy H1, BioTek) and performed with excitation and emission wavelengths at 485 and 528 nm, respectively. Depending on the standard curve of “sfGFP (µg / mL) - Fluorescence (a.u.)” (**Figure S11**, **Supporting Information**), the sfGFP yield was calculated as µg / mL.

### Lyophilization and Rehydration of EcN CFPS Mixture

Before lyophilization, each 15 µL CFPS mixture (without plasmid) was prepared and loaded into 1.5 mL tube and performed quick-freezing by liquid nitrogen. Then, the tubes were lyophilized by LABCONCO FreeZone at −70 °C, 70 Pa for at least 12 h. Next, the lyophilized CFPS mixture was stored at room temperature (25 °C) to be used. When rehydrate CFPS mixture, 15 µL nuclease free water (with plasmid) was added to tube and incubated at 30 °C for at least 3 h.

## Author Information

### Author Contributions

J.L. and F.B. designed the experiments. F.B. performed all experiments. Y.Z. helped perform molecular cloning. X.J. helped prepare cell extracts. F.B. analyzed the data and drafted the manuscript. J.L., W.-Q.L. and S.L. revised and edited the manuscript. J.L. conceived and supervised the study. All authors read and approved the final manuscript.

## Supporting information

Plasmid Sequence

## Acknowledgments

This work was supported by grants from the National Natural Science Foundation of China (31971348 and 32171427).

