## Supporting Information for "Expanding the toolbox of probiotic *Escherichia coli* Nissle 1917 for synthetic biology"

I. Supporting Tables

I. Supporting Tables

### Supporting Table S1. *E. coli* strains.

| Strain name | Purpose | Genotype | Origin |
|---|---|---|---|
| Mach1-T1 | Molecular cloning | $F^-$ φ80(*lac*Z)Δ*M15* Δ*lac*X74 *hsd*R($r_K^-$ $m_K^+$) Δ*rec*A1398 *end*A1 *ton*A | TransGen Biotech |
| MG1655 | Type strain | K12 $F^-$ *lambda- ilvG- rfb-50 rph-1* | Shanghai Weidi Biotechnology |
| S17-1 λpir | Cloning R6K-ori plasmids, donor cell for conjugation transfer of “RP4-oriT” derived plasmids | *RP4-2 (Km::Tn7,Tc::Mu-1)*, *pro*-82, *LAMpir*, *rec*A1, *end*A1, *thi*E1, *hsd*R17, *cre*C510 | Shanghai Weidi Biotechnology |
| DH5α λpir | Cloning R6K-ori plasmids | $F^-$ φ80 *lac*ZΔM15 Δ(*lac*ZYA-*arg* F) *LAMpir* U169 *end*A1 *rec*A1 *hsd*R17($r_k^-$,$m_k^+$) *sup*E44λ- *thi* -1 *gyr*A96 *rel*A1 *pho*A | Shanghai Weidi Biotechnology |
| Nissle 1917 (EcN) | Probiotic | | Mutaflor |

### Supporting Table S2. Genetic parts.

| Part name | Type | DNA sequence | Reference |
|---|---|---|---|
| oriV (ColE2) | origin of replication | acaaaacaacctatatcagataacag | This work |
| BioBrick prefix | BioBrick assembly site | gaattcgcggccgcttctagag | 1 |
| BioBrick suffix | BioBrick assembly site | tactagtagcggccgctgcag | 1 |
| J23100 | promoter | ttgacggctagctcagtcctaggtacagtgctagc | 1, BBa_J23100 |
| J23105 | promoter | tttacggctagctcagtcctaggtactatgctagc | 1, BBa_J23105 |
| J23106 | promoter | tttacggctagctcagtcctaggtatagtgctagc | 1, BBa_J23106 |
| J23109 | promoter | tttacagctagctcagtcctagggactgtgctagc | 1, BBa_J23109 |
| J23113 | promoter | ctgatggctagctcagtcctagggattatgctagc | 1, BBa_J23113 |
| J23114 | promoter | tttatggctagctcagtcctaggtacaatgctagc | 1, BBa_J23114 |
| paraBAD | promoter | aaagccatgacaaaaacgcgtaacaaaagtgtctataatcacggcagaaaagtccacattgattatttgcacggcgtcacactttgctatgccatagcatttttatccataagattagcggatcctacctgacgctttttatcgcaactctctactgtttctccat | From *E. coli* MG1655 genome (GenBank: U00096.3) |
| pT7 | promoter | taatacgactcactataggg | From plasmid vector pET22b |
| B0034 | RBS | aaagaggagaaa | 1, BBa_B0034 |
| B0031 | RBS | tcacacaggaaacc | 1, BBa_B0031 |
| B0032 | RBS | tcacacaggaaag | 1, BBa_B0032 |
| B0033 | RBS | tcacacaggac | 1, BBa_B0033 |
| B0015 | terminator | ccaggcatcaaataaaacgaaaggctcagtcgaaagactgggcctttcgttttatctgttgtttgtcggtgaacgctctctactagagtcacactggctcaccttcgggtgggcctttctgcgtttata | 1, BBa_B0015 |
| T7 terminator | terminator | ctagcataaccccttggggcctctaaacgggtcttgaggggttttttg | From plasmid vector pET22b |
| *attP* (int 1) | integrase recognition site (**core sequence**) | ttattgccagtcagatggggcatgcgagcgcgcagatggtgttcaatgtttacggtgcatggatggctgacagcagcgcagagcagatcgcaatgctgaatcagaagctggcagattttgccccattgatgccccatagccacgagaacagtacgggaggattattaaaatcagtaagttaacccctaacgcccgtcatg**ttaactg**tgtgacgcgggcattttaaaaatcactaaagaacgcccaagagcatgtgttttctttagtttattcaatgcattaaaaaatagtttcgcatgaaattcggtaaacttcatgtgtgcaataatgtcccattcatgccccaaaatgccccaaagcagacatttttgccccaagtatgccccacaagtcacgtcttcaagtcgtcta | This work |
| *attB* (int 1) | integrase recognition site (**core sequence**) | tgattgccgaagatgcctgtagcgccgctagcgccgagcagcacaataacagcattaatcatatctacccgcgcatcgcccgtgtgcgtagcgttgaagagatcctcaacgcgttatgatttacatcggtttgccacaatggtcgcatcctaaatgggtgcggttggggatcaccagccttgaagagtatgcccgccact**ttaactg**cgtggagggtaacaccacgctttatgccctgccgaaacccgaggttgtcctgcgctggcgtgagcagaccacagatgacttccgcttctgttttaagtttccggcgaccatttcgcatcaggcagcattacggcattgcgatgatttagtgactgaatttttgacccgcatgtcaccgttggctccgcgcattggacaatactg | This work |
| *attP* (int 2) | integrase recognition site (**core sequence**) | atcagagacgggggaatggctaagttgttgttaagagtgtgaactcgagagcataaaagtggatgcccgacgcgcggcaaaatggtaaaaaacagcatcagtagtaagaaaaattgagcgagagaaaggaattttaatgcagggatttatgcatgacaaaatcaaaaatgctttgtaagtgattgattttgttgaatgat**tggcggagagaggg**tctaaatttttagtgttcgacaatgcccaccaaagtgctatttttcctttaaaaacaaaatattgtcattatttttttgttcggtagtgtattgctactgtgcaaacaatcagctattttatgcagtaccgtatgtatggcgggttgtatgacaggattttgctatgccaaagaaggccaaggaactctcaggactcgtc | This work |
| *attB* (int 2) | integrase recognition site | taatggtgaccgataagaaaacgactgaataactgtagattttcgctcgaaaccttcctgtcagacccata | This work |

| | | | |
|---|---|---|---|
| | (**core sequence**) | gcgaatcaagtgctgaatgtcacagtatcgaacagaaaacagtgacgatctaacccttcaagaatattctacgattgttctgtttaggaaaagcaaggcgggaagtcgggagataagtcattgataaag**tggcggagagaggg**ggatttgaaccccccggtagagttgcccctactccggttttcgagaccggtccgttcagccgctccggcatctctccgttcagatggttgccatgatgccaggaaatttggcattttaacagtccctgtccgtgcaattttgttcaagtgacgagtttgcgagcaaaacgatgattaagtggccctggaaagtacaagaat | |
| *attP* (int 3) | integrase recognition site (**core sequence**) | gctctttagacacagttccactaggggagaggatcagcgcttcccacgtagggacactatagtaactatcaccacatgcagtagggaagctcaaaccaaatttatgtagccagctcctattggtggtcattctggtggtcttgacaggaagataactctggtttagcttacttattagccacttactggcaaaggcgatc**ccagtcagaggagccaa**atttcctgctttcatgcatccttgcgaatccttatgttatttttattcaacaggttagcgtgaaaactcttcccggtgcattttgattttaccctctgcatcgggaaaaattggtggtcaaatctggggtcaggttagttcgataatggagtgacccccatatgtcccttaacgacgcaaaaatccgtagtctcaagccc | This work |
| *attB* (int 3) | integrase recognition site (**core sequence**) | gccgcaaaagtgtgaaaaaaagccagttgtatcggaataaacgacaaaatgcagattatttcagcaaacgatttcaaatttaaaaaacaggctttgacattgtgggtgggcatcgctaatattcgcctcgttctcacgattcctctgtagttcagtcggtagaacggcggactgttaatccgtatgtcactggttcgagt**ccagtcagaggagccaa**ttcaaaaaagcctgctttcgagcaggctttttactttctaattaccaacgctcttaaaacatctgtcttgaaccagaactaatttgcacaggcattcccgatcgatgttgcaacgcagcatttgcgcgatttacatcaacttcttgcccgttgataaacgcccgcaaagatggggttaccggcaatggcacttttcggtc | This work |
| *attP* (int 4) | integrase recognition site (**core sequence**) | ctctgaatggtggaaatcaccacgcagatcatccaccagcttcgtttttataaattaaacatccgggaaaatccgaccctgaattgtgcatttttcagcaaatcattttagttatacgtttggttatacggtttaaaagttgaatcatatacattattttaaatcatgaagttacacgctcaattaaactactctcgtg**gaccaaa**tcgcccaccggatatcaccggaaagcaccggataagacagggataacccgctaatcacttgattggcgggtttttttatgtttttttagtcaccgaacatcaccggatttataagctatcagccggatttttttagttatacgtttagttatacgggctgacgtataactaaagccgtataactaaaaccctgtttttcgtg | This work |
| *attB* (int 4) | integrase recognition site (**core sequence**) | ttaagtttgttattgcaagtactgacagacgagaagcgttttatcgctaactgattaattataaatcagttagcaaaatatcttacttgcaatcggtgtggaaaacggtagtattagcagccacgagtcggcacgtagcgcagcctggtagcgcaccgtcatggggtgtcgggggtcggaggttcaaatcctctcgtgcc**gaccaaa**aatcccaagaaaaccaacctattgcggttggtttttttatatctgcatttaattcgataaacagacagcgacacatcacggcctgtttattttctgttatcagaacgtccagactacacccgcctgagttgtgctgctttcttcacgggagaaacggctgtcgataccaagcgttagccccgtcaatgtttgctcaaagc | This work |
| sfGFP | CDS | atgcgtaaaggcgaagagctgttcactggtgtcgtccctattctggtggaactggatggtgatgtcaacggtcataagttttccgtgcgtggcgagggtgaaggtgacgcaactaatggtaaactgacgctgaagttcatctgtactactggtaaactgccggtaccttggccgactctggtaacgacgctgacttatggtgttcagtgctttgctcgttatccggaccatatgaagcagcatgacttcttcaagtccgccatgccggaaggctatgtgcaggaacgcacgatttcctttaaggatgacggcacgtacaaaacgcgtgcggaagtgaaatttgaaggcgataccctggtaaaccgcattgagctgaaaggcattgactttaaagaagacggcaatatcctgggccataagctggaatacaattttaacagccacaatgtttacatcaccgccgataaacaaaaaaatggcattaaagcgaattttaaaattcgccacaacgtggaggatggcagcgtgcagctggctgatcactaccagcaaaacactccaatcggtgatggtcctgttctgctgccagacaatcactatctgagctaccaaagcgttctgtctaaagatccgaacgagaaacgcgatcatatggttctgctggagttcgtaaccgcagcgggcatcacgcatggtatggatgaactgtactaa | 2 |
| araC | CDS | atggctgaagcgcaaaatgatcccctgctgccgggatactcgtttaatgcccatctggtggcgggtttaacgccgattgaggccaacggttatctcgatttttttatcgaccgaccgctgggaatgaaaggttatattctcaatctcaccattcgcggtcaggggggtggtgaaaaatcagggacgagaatttgtttgccgaccgggtgatattttgctgttcccgccaggagagattcatcactacggtcgtcatccggaggctcgcgaatggtatcaccagtgggtttactttcgtccgcgcgcctactggcatgaatggcttaactggccgtcaatatttgccaatacggggttctttcgcccggatgaagcgcaccagccgcatttcagcgacctgtttgggcaaatcattaacgccgggcaaggggaa | From *E. coli* MG1655 genome (GenBank: U00096.3) |

| | | | |
|---|---|---|---|
| | | gggcgctattcggagctgctggcgataaatctgcttgagcaattgttactgcggcgcatggaagcgattaacgagtcgctccatccaccgatggataatcgggtacgcgaggcttgtcagtacatcagcgatcacctggcagacagcaattttgatatcgccagcgtcgcacagcatgtttgcttgtcgccgtcgcgtctgtcacatcttttccgccagcagttagggattagcgtcttaagctggcgcgaggaccaacgtatcagccaggcgaagctgcttttgagcaccacccggatgcctatcgccaccgtcggtcgcaatgttggttttgacgatcaactctatttctcgcgggtatttaaaaaatgcaccggggccagcccgagcgagttccgtgccggttgtgaagaaaaagtgaatgatgtagccgtcaagttgtcataa | |
| T7 RNA polymerase | CDS | atgaacacgattaacatcgctaagaacgacttctctgacatcgaactggctgctatcccgttcaacactctggctgaccattacggtgagcgtttagctcgcgaacagttggcccttgagcatgagtcttacgagatgggtgaagcacgcttccgcaagatgtttgagcgtcaacttaaagctggtgaggttgcggataacgctgccgccaagcctctcatcactaccctactccctaagatgattgcacgcatcaacgactggtttgaggaagtgaaagctaagcgcggcaagcgcccgacagccttccagttcctgcaagaaatcaagccggaagccgtagcgtacatcaccattaagaccactctggcttgcctaaccagtgctgacaatacaaccgttcaggctgtagcaagcgcaatcggtcgggccattgaggacgaggctcgcttcggtcgtatccgtgaccttgaagctaagcacttcaagaaaaacgttgaggaacaactcaacaagcgcgtagggcacgtctacaagaaagcatttatgcaagttgtcgaggctgacatgctctctaagggtctactcggtggcgaggcgtggtcttcgtggcataaggaagactctattcatgtaggagtacgctgcatcgagatgctcattgagtcaaccggaatggttagcttacaccgccaaaatgctggcgtagtaggtcaagactctgagactatcgaactcgcacctgaatacgctgaggctatcgcaacccgtgcaggtgcgctggctggcatctctccgatgttccaaccttgcgtagttcctcctaagccgtggactggcattactggtggtggctattgggctaacggtcgtcgtcctctggcgctggtgcgtactcacagtaagaaagcactgatgcgctacgaagacgtttacatgcctgaggtgtacaaagcgattaacattgcgcaaaacaccgcatggaaaatcaacaagaaagtcctagcggtcgccaacgtaatcaccaagtggaagcattgtccggtcgaggacatccctgcgattgagcgtgaagaactcccgatgaaaccggaagacatcgacatgaatcctgaggctctcaccgcgtggaaacgtgctgccgctgctgtgtaccgcaaggacaaggctcgcaagtctcgccgtatcagccttgagttcatgcttgagcaagccaataagtttgctaaccataaggccatctggttcccttacaacatggactggcgcggtcgtgtttacgctgtgtcaatgttcaacccgcaaggtaacgatatgaccaaaggactgcttacgctggcgaaaggtaaaccaatcggtaaggaaggttactactggctgaaaatccacggtgcaaactgtgcgggtgtcgataaggttccgttccctgagcgcatcaagttcattgaggaaaaccacgagaacatcatggcttgcgctaagtctccactggagaacacttggtgggctgagcaagattctccgttctgcttccttgcgttctgctttgagtacgctggggtacagcaccacggcctgagctataactgctcccttccgctggcgtttgacgggtcttgctctggcatccagcacttctccgcgatgctccgagatgaggtaggtggtcgcgcggttaacttgcttcctagtgaaaccgttcaggacatctacgggattgttgctaagaaagtcaacgagattctacaagcagacgcaatcaatgggaccgataacgaagtagttaccgtgaccgatgagaacactggtgaaatctctgagaaagtcaagctgggcactaaggcactggctggtcaatggctggcttacggtgttactcgcagtgtgactaagcgttcagtcatgacgctggcttacgggtccaaagagttcggcttccgtcaacaagtgctggaagataccattcagccagctattgattccggcaagggtctgatgttcactcagccgaatcaggctgctggatacatggctaagctgatttgggaatctgtgagcgtgacggtggtagctgcggttgaagcaatgaactggcttaagtctgctgctaagctgctggctgctgaggtcaaagataagaagactggagagattcttcgcaagcgttgcgctgtgcattgggtaactcctgatggtttccctgtgtggcaggaatacaagaagcctattcagacgcgcttgaacctgatgttcctcggtcagttccgcttacagcctaccattaacaccaacaaagatagcgagattgatgcacacaaacaggagtctggtatcgctcctaactttgtacacagccaagacggtagccaccttcgtaagactgtagtgtgggcacacgagaagtacggaatcgaatcttttgcactgattcacgactccttcggtaccattccggctgacgctgcgaacctgttcaaagcagtgcgcgaaactatggttgacacatatgagtcttgtgatgtactggctgatttctacgaccagttcgctgaccagttgcacgagtctcaattggacaaaatgccagcacttccggctaaaggtaacttgaacctccgtgacatcttagagtcggacttcgcgttcgcgtaa | From *E. coli* BL21 (DE3) genome (GenBank: CP001509.3) |
| lacI | CDS | gtgaaaccagtaacgttatacgatgtcgcagagtatgccggtgtctcttatcagaccgtttcccgcgtggtga | From *E. coli* |

| | | | |
|---|---|---|---|
| | | accaggccagccacgtttctgcgaaaacgcgggaaaaagtggaagcggcgatggcggagctgaattacattcccaaccgcgtggcacaacaactggcgggcaaacagtcgttgctgattggcgttgccacctccagtctggccctgcacgcgccgtcgcaaattgtcgcggcgattaaatctcgcgccgatcaactgggtgccagcgtggtggtgtcgatggtagaacgaagcggcgtcgaagcctgtaaagcggcggtgcacaatcttctcgcgcaacgcgtcagtgggctgatcattaactatccgctggatgaccaggatgccattgctgtggaagctgcctgcactaatgttccggcgttatttcttgatgtctctgaccagacacccatcaacagtattattttctcccatgaagacggtacgcgactgggcgtggagcatctggtcgcattgggtcaccagcaaatcgcgctgttagcgggcccattaagttctgtctcggcgcgtctgcgtctggctggctggcataaatatctcactcgcaatcaaattcagccgatagcggaacgggaaggcgactggagtgccatgtccggttttcaacaaaccatgcaaatgctgaatgagggcatcgttcccactgcgatgctggttgccaacgatcagatggcgctgggcgcaatgcgcgccattaccgagtccgggctgcgcgttggtgcggatatctcggtagtgggatacgacgataccgaagacagctcatgttatatcccgccgttaaccaccatcaaacaggattttcgcctgctggggcaaaccagcgtggaccgcttgctgcaactctctcagggccaggcggtgaagggcaatcagctgttgcccgtctcactggtgaaaagaaaaaccaccctggcgcccaatacgcaaaccgcctctccccgcgcgttggccgattcattaatgcagctggcacgacaggtttcccgactggaaagcgggcagtga | MG1655 genome (GenBank: U00096.3) |
| int 1 | CDS | atggataaagtcacatatccaacaggcgtcgaaaaccacggtggcacattacgcatctggtttaattttaaaggtaagcgtgtcagggaaagtctcggtgtccctgacaccgctaagaacaggaagatagccggggaactgcggacatcagtatgttttgccatccgcacaggaacctttgattatgcaacccagtttcctgactcccctaacctcaaggcttttggtgtaagtaaaaaagacattacagtgaaagaacttgaagaaaaatggctggatctgaaacggatggaaatctgcgcgaacgcatttaatcgctatgaatctgtcgcaaggaatatggtgccgaggatcggaggtaatcgcctggtgtcagcagtaaccaaagaggaattgctgtatctgaggaaatatttgctaactggttatcagaatccgacgaaaaacaaagccccggcaaaagggcgaagcgttgttactgtgaactattacatgacgacaatggccggaatgtttcagtttgctgcggatcacggttacttagaggtgaacccattcgagggaattaagcctctgaaaaaagccagggcagaaccagatcctctgtctcgtgatgaatttattcgcctgatagatgcatgccggcatcagcagacgaaaaacctgtggtcattagcagtgtacacaggaatgcgtcacggggaactggtctccctggcctgggaagatatcgacctgaaggctggaacaattaccgtcagacgtaattatacgaaacttggtgagttcactctaccgaaaaccgaggcaagcacagatcgagtggtgcatcttatccagcccgcaatcagtatcctgaaaaatcaggctgaaatgacaaggctgggcaggcaatatcacattgaagtgcagttacgtgagtacggccgttcggtgaaccatgagtgtacattcgtctttaatccgcatgtggtcagacgcagtaagcaggtcggatttatctaccgggtcgattcagtaggcgactcatgggaagcggcacttaagcgtgcggggatcagacacagaaaggcgtaccagtcacgacacacctatgcgtgctggtcattatcagctggtgcaaaccctagttttattgccagtcagatggggcatgcgagcgcgcagatggtgttcaatgtttacggtgcatggatggctgacagcagcgcagagcagatcgcaatgctgaatcagaagctggcagattttgccccattgatgccccatagccacgagaacagtacgggaggattattaaaatcagtaagttaa | This work |
| int 2 | CDS | atgccaaagaaggccaaggaactctcaggactcgtcgtatcacgattaaaatccgaaggcatgtatgccgttggtggtgtggatggtttataccttcgcatccgaaatcagtcccgtgcatgggttctttgcgtggctatgggaacccgaatcaacaatctcggtagaacagttccaagacgtttgaatatgggtcttggcccttacccggaagtctccttagccgaagcgcgtgacaaagcgcgtgagctacgtaaacaaatccgtaatggtattaatccccttcaggaaaaacacgagcaaaaagcccggcaggaaatactggcccggaagaaaaagacctttgccgaatgttgtgaagaagtgctggaagtcaaagacagtgagatgaagaacaaaaagcatctcgcgcaatggcgttccacactggagacctatgcctaccctttcattggcaaaaaggccgttagtgaaatcaccaaagtcgatcttctggcaatactggaacctatctggttaaccaagaatgaaaccgccagtcgtctacgtggacgtattgaaacggtgatcgattacgccaaggccaaagaatactttgaaggtgataatcctgctgcatggaaaggtatgctgaaacctctcctgcctcagccaagtaaagttcagatcaccaaacaccatgcggctcttccctataaccagatcggctcctttatgaaagaactgcgtgaacgaagcggcgtttcgcctcgtgcgcttgagtttgccatactgaccgcagcacgctcaggtgaaattcgtggtgccgagtggtcagagatcgatcttgaaggtaaaacatggac | This work |

| | | | |
|---|---|---|---|
| | | catccccgccagtcgaatgaaagcaacgaaggagcatcgagttccgttgtctgatgctgctgttgccctgtt<br>aaaggctttaccacgctttaaaggtatcaattttgtattccctgccactcggaagggacaactctcggatact<br>gctttactggcagtcctaaagcgaatgggatataccgacttaacgcagcatgggtttagatcgacattccgt<br>gattgggctggtgaaacgaccaactacccgcgtgaggtaattgaacatgcgttagctcatcagttggcaa<br>ataaggcagaagctgcgtatcaacgtgggacgttatggcctaagcgggtggcgttaatggatgattgggc<br>ggggtattgtatttcttag | |
| int 3 | CDS | atgtcccttaacgacgcaaaaatccgtagtctcaagcccactgataaacccttaaagtctccgattcccac<br>ggtctgtatctgctggtcaaaccaggcggttcccgtctctggtatcttaaataccgtattaatggcaaagaatc<br>ccgtattgcactgggtgcctaccctgccgtctccctgtctgacgcccgccagcagcgcgaaggcatccgta<br>agatgctggcgctgaacatcaacccggcgcagcaacgtgccgctgaacgcggctcgcgcatgcagga<br>gaaaatgtttaaatccgtggcactggagtggcacagtagcaagaaaaaatggtcgcagaataccgcag<br>atcgcgtgcttgcccgtctgaacagacatgtctttccgaccatcgggcatctaccagttaccgagcttaaatc<br>acgtcatttcattgaattactgaaaggcattgaggaaaagggcttgcttgaagttgcgtcccgctcgcggca<br>acacctcagcaacattatgcgttatgctgttcatcaggggttaatcgaaatcaacccagctgcaaatcttga<br>cggcgtgacggcttctcccgccagacgacattatcccaccctgccactggagcgccttcctgagctgcttg<br>aacggattgacagttatcaccaaggacgggaactcaccaggcttgcagtcctgctgacgttgcatgtgttc<br>attcgctcaagtgaactgcgctatgctcgttggacagagattaacttcagaaatcggatctggacgatacct<br>gccacacgcgaagcgattgcaggcgtacgttattccagtcgaggtgccaaaatgcgtacaccgcatattg<br>tgcccctttctgagcaggttatctctattctgaaaaggattaaggaaatatctggcgggtatgaactggtgttc<br>cccggctatcatgacccctacaagccgatgagtgaaaatactatcaataaggcacttcgccagatgggat<br>acaacacgaaacaagatatctgcggtcatggtttccgggcaatggcgtgtagcgcgctgatggaatccg<br>gtttatggtcgcaagatgcagtggagcgtcagatgagtcaccaagagcgcaacactgtgcgtctggctta<br>cattcataaggccgaacatatggaagcccgcatggacatgatgcagtggtggtcggattatctggatatgt<br>gcagtgaaatctgggtgcccccgtacatctggagtcagcaaaacattaatctcgcagtaacctga | This work |
| int 4 | CDS | atggcgagaaaaacacacccattaaccacagtgcagatcaaagcagccagaccagcgcaaaagga<br>gtacaccctgcaggacggcggagggctttttctcctggtcaaaccgtctggatcaaaactctggagattttc<br>ctactaccgaccatcggacaaaaaaagaatattgctgagttttggatcgcttgaagatgtttccctggctgat<br>gccagaaaacgccgtagcgagtacaggacgttaatcagtgccggaactgacccgcaggaccacgag<br>aggcaaaaaagagagacagaggcccgaagacaagggaacacgttcgaaaatgtggcggcggcatg<br>gtaccaggtgaaaatcagccagaatctggcccccaacacgattaaagacatctggcgttcgctggataa<br>atatgtattcccgttcatcggcaacacgccaatagataccctcaccgcccgaaggttcgttgaagtgcttac<br>acccatcaaggagcgcggcaacctggaaacactcaagcgggttttacagcgcgttaatgaggtaatgg<br>attacgccgccaacagtgggctgattgatgccaatccggctatgaatgtgcgtaaggcgtttccctcacctg<br>taaaaaaacatatgccaacaatccgccccgaacagctgccggagcttatgcaggctttatcagtatcggc<br>aacagaacggcagaccagattactgattgaatggcagttactgaccgtaacccgtcccgccgaagcgtc<br>atcaacgcggtgggatgaaatcaacctggacgcgaagcaatggacgatacctgccggacgcatgaag<br>atgcgcagggatcacgttatcccgctttccggtcaggctatggcggtgctggaggccatgaaaccaatca<br>gccaccaccgcaattacgttttcccaagtctgaaagacccacagcagccgatgaacagccagacagct<br>aacgcagcattgcggcgtatgggattcgctggcgtgctggtgtctcatggattacgcgccatattcagcaca<br>gcagcgaacgaggaaggattcgagccggacgtaatagaggcggcacttgcccacgtcgacaccaac<br>gaagttagacgggcatacaaccggagcaactacatagaaaaacgcatcgtgctgatgcgctggtgggg<br>cgaatttgtcgaggctgcggcgacgggcgtaaccctcgccagtggtaaaaggggtatccgagccgtgta<br>g | This work |
| sfGFP (CFPS) | CDS | atgagcaaaggtgaagaactgtttaccggcgttgtgccgattctggtggaactggatggcgatgtgaacg<br>gtcacaaattcagcgtgcgtggtgaaggtgaaggcgatgccacgattggcaaactgacgctgaaatttat<br>ctgcaccaccggcaaactgccggtgccgtggccgacgctggtgaccaccctgacctatggcgttcagtgt | From plasmid pJL1-sfGFP (Addgene: |

| | | | |
|---|---|---|---|
| | | tttagtcgctatccggatcacatgaaacgtcacgatttctttaaatctgcaatgccggaaggctatgtgcaggaacgtacgattagctttaaagatgatggcaaatataaaacgcgcgccgttgtgaaatttgaaggcgatacctggtgaaccgcattgaactgaaaggcacggattttaaagaagatggcaatatcctgggccataaactggaatacaactttaatagccataatgtttatattacggcggataaacagaaaaatggcatcaaagcgaattttaccgttcgccataacgttgaagatggcagtgtgcagctggcagatcattatcagcagaataccccgattggtgatggtccggtgctgctgccggataatcattatctgagcacgcagaccgttctgtctaaagatccgaacgaaaaaggcacgcgggaccacatggttctgcacgaatatgtgaatgcggcaggtattacgtggagccatccgcagttcgaaaaataa | 102634) |
| lacO (lac operator) | operator (transcriptional repressor DNA-binding site) | ggaattgtgagcggataacaattcc | From plasmid vector pET22b |
| RP4-oriT | oriT | gaataagggacagtgaagaaggaacacccgctcgcgggtgggcctacttcacctatcctgcccggctgacgccgttggatacaccaaggaaagtctaca | 1, BBa_K125320 (GenBank: X14165.1) |
| tracrRNA scaffold | tracrRNA scaffold for spCas9 | gttttagagctagaaatagcaagttaaaataaggctagtccgttatcaacttgaaaaagtggcaccgagtcggtgcttttttt | 3 |

**Supporting Table S3. Vectors.**

| Vector name | Antibiotic resistance gene | Origin of replication | Reference |
|---|---|---|---|
| pMUT1 | N/A | ColE1 | GenBank: CP058218.1 |
| pMUT2 | N/A | ColE2 | GenBank: CP058219.1 |
| pFB1 | AmpR | p15A | 2 |
| pSB1C3 | CmR | ColE1 | 1 (pSB1C3), 2 |
| pFB2 | KanR | pSC101 | 2 |
| pFB3 | SmR | incW | 2 |
| pFB4 | CmR | p15A | This work |
| pET22b | AmpR | ColE1+rop | This work |
| pYF1 | KanR | R6K | This work |
| pKD4 | AmpR | R6K | 4 |
| pKD46 | AmpR | pSC101 | 4 |
| pFB5 | SmR | R6K | This work |
| pJL1 | KanR | ColE1 | Addgene: 102634 |

**Supporting Table S4. Plasmids.**

| Plasmid name | Biobricks | Vector | Antibiotic resistance gene | Origin of replication |
|---|---|---|---|---|
| pFB341 | pMUT1-tru | pMUT1 | N/A | ColE1 |
| pFB342 | pMUT1-mini (with BioBrick prefix / suffix) | pMUT1 | CmR | ColE1 |
| pFB343 | pMUT2-tru | pMUT2 | N/A | ColE2 |
| pFB344 | pMUT2-mini (with BioBrick prefix / suffix) | pMUT2 | KanR | ColE2 |
| pFB345 | RP4-oriT | pFB1 | AmpR | p15A |
| pFB346 | J23100 - B0034 - sfGFP - (6xHis) - B0015 | pSB1C3 | CmR | ColE1 |
| pFB347 | RP4-oriT | pSB1C3 | CmR | ColE1 |
| pFB348 | J23100 - B0034 - sfGFP - (6xHis) - B0015 | pFB1 | AmpR | p15A |
| pFB349 | J23105 - B0034 - sfGFP - (6xHis) - B0015 | pSB1C3 | CmR | ColE1 |
| pFB350 | J23106 - B0034 - sfGFP - (6xHis) - B0015 | pSB1C3 | CmR | ColE1 |
| pFB351 | J23109 - B0034 - sfGFP - (6xHis) - B0015 | pSB1C3 | CmR | ColE1 |
| pFB352 | J23113 - B0034 - sfGFP - (6xHis) - B0015 | pSB1C3 | CmR | ColE1 |
| pFB353 | J23114 - B0034 - sfGFP - (6xHis) - B0015 | pSB1C3 | CmR | ColE1 |
| pFB354 | araC - paraBAD - B0034 - sfGFP - (6xHis) - B0015 | pSB1C3 | CmR | ColE1 |
| pFB355 | J23113 - B0031 - T7 RNA polymerase - B0015 | pFB4 | CmR | p15A |
| pFB356 | J23113 - B0032 - T7 RNA polymerase - B0015 | pFB4 | CmR | p15A |
| pFB357 | J23113 - B0033 - T7 RNA polymerase - B0015 | pFB4 | CmR | p15A |
| pFB358 | J23113 - B0034 - T7 RNA polymerase - B0015 | pFB4 | CmR | p15A |
| pFB359 | lacI - pT7 - lacO - sfGFP - (6xHis) - T7 terminator | pET22b | AmpR | ColE1+rop |
| pFB360 | araC - paraBAD - B0034 - (6xHis) - int 1 - B0015 | pSB1C3 | CmR | ColE1 |
| pFB361 | araC - paraBAD - B0034 - (6xHis) - int 2 - B0015 | pSB1C3 | CmR | ColE1 |
| pFB362 | araC - paraBAD - B0034 - (6xHis) - int 3 - B0015 | pSB1C3 | CmR | ColE1 |
| pFB363 | araC - paraBAD - B0034 - (6xHis) - int 4 - B0015 | pSB1C3 | CmR | ColE1 |
| pFB364 | int 1: *attP* (*attPL* + *attPR*) - *attB* (*attBL* + *attBR*) | pFB1 | AmpR | p15A |
| pFB365 | int 2: *attP* (*attPL* + *attPR*) - *attB* (*attBL* + *attBR*) | pFB1 | AmpR | p15A |
| pFB366 | int 3: *attP* (*attPL* + *attPR*) - *attB* (*attBL* + *attBR*) | pFB1 | AmpR | p15A |
| pFB367 | int 4: *attP* (*attPL* + *attPR*) - *attB* (*attBL* + *attBR*) | pFB1 | AmpR | p15A |
| pFB368 | int 1: *attP* (*attPL* + *attPR*) - J23100 - B0034 - sfGFP - (6xHis) - B0015 - *attB* (*attBL* + *attBR*) | pFB1 | AmpR | p15A |
| pFB369 | int 2: *attP* (*attPL* + *attPR*) - J23100 - B0034 - sfGFP - (6xHis) - B0015 - *attB* (*attBL* + *attBR*) | pFB1 | AmpR | p15A |
| pFB370 | int 3: *attP* (*attPL* + *attPR*) - J23100 - B0034 - sfGFP - (6xHis) - B0015 - *attB* (*attBL* + *attBR*) | pFB1 | AmpR | p15A |
| pFB371 | int 4: *attP* (*attPL* + *attPR*) - J23100 - B0034 - sfGFP - (6xHis) - B0015 - *attB* (*attBL* + *attBR*) | pFB1 | AmpR | p15A |
| pFB372 | int 1: *attP* (*attPL* + *attPR*) - J23100 - B0034 - sfGFP - (6xHis) - B0015 - *attB* (*attBR* + *attBL* as inversion) | pFB1 | AmpR | p15A |
| pFB373 | int 2: *attP* (*attPL* + *attPR*) - J23100 - B0034 - sfGFP - (6xHis) - B0015 - *attB* (*attBR* + *attBL* as inversion) | pFB1 | AmpR | p15A |
| pFB374 | int 3: *attP* (*attPL* + *attPR*) - J23100 - B0034 - sfGFP - (6xHis) - B0015 - *attB* (*attBR* + *attBL* as inversion) | pFB1 | AmpR | p15A |

| | | | | |
|---|---|---|---|---|
| pFB375 | int 4: *attP* (*attPL* + *attPR*) - J23100 - B0034 - sfGFP - (6xHis) - B0015 - *attB* (*attBR* + *attBL* as inversion) | pFB1 | AmpR | p15A |
| pFB376 | int 1 *attB* - int 2 *attB* - int 3 *attB* - int 4 *attB* | pSB1C3 | CmR | ColE1 |
| pFB377 | (pYF1) - *attP*-TP901-1 - int 1 *attB* - int 2 *attB* - int 3 *attB* - int 4 *attB* | pYF1 | KanR | R6K |
| pFB378 | (derived from pKD4) homoL (csgA) - *attB*-TP901-1 - FRT - KanR - FRT - homoR (csgA) | pKD4 | AmpR, KanR | R6K |
| pFB379 | araC - paraBAD - TP901-1 - (6xHis) - T7 terminator | pKD46 | AmpR | pSC101 |
| pFB380 | int 1: *attP* | pFB5 | SmR | R6K |
| pFB381 | int 2: *attP* | pFB5 | SmR | R6K |
| pFB382 | int 3: *attP* | pFB5 | SmR | R6K |
| pFB383 | int 4: *attP* | pFB5 | SmR | R6K |
| pFB384 | J23105 - B0034 - T7 RNA polymerase - B0015 | pFB1 | AmpR | p15A |
| pFB385 | J23106 - B0034 - T7 RNA polymerase - B0015 | pFB1 | AmpR | p15A |
| pFB386 | J23109 - B0034 - T7 RNA polymerase - B0015 | pFB1 | AmpR | p15A |
| pFB387 | J23113 - B0034 - T7 RNA polymerase - B0015 | pFB1 | AmpR | p15A |
| pFB388 | J23114 - B0034 - T7 RNA polymerase - B0015 | pFB1 | AmpR | p15A |
| pFB389 | pT7 - RBS - sfGFP (CFPS) | pJL1 | KanR | ColE1 |
| pFB390 | J23100 - crRNA-p15A-1 - tracrRNA scaffold | pSB1C3 | CmR | ColE1 |
| pFB391 | J23100 - crRNA-p15A-2 - tracrRNA scaffold | pSB1C3 | CmR | ColE1 |
| pFB392 | J23100 - crRNA-p15A-3 - tracrRNA scaffold | pSB1C3 | CmR | ColE1 |
| pFB393 | J23100 - crRNA-incW-1 - tracrRNA scaffold | pSB1C3 | CmR | ColE1 |
| pFB394 | J23100 - crRNA-incW-2 - tracrRNA scaffold | pSB1C3 | CmR | ColE1 |
| pFB395 | J23100 - crRNA-incW-3 - tracrRNA scaffold | pSB1C3 | CmR | ColE1 |

**Supporting Table S5. DNA sequences for pCas system-based crRNA transcription.**

| Name | Sequence (5' to 3') | Purpose | Reference |
|---|---|---|---|
| crRNA-p15A-1 | gctctcctgttcctgccttt | Curing p15A-oriV derived plasmids | This work |
| crRNA-p15A-2 | ctatcgtcttgagtccaacc | Curing p15A-oriV derived plasmids | This work |
| crRNA-p15A-3 | gttacctcggttcaaagagt | Curing p15A-oriV derived plasmids | This work |
| crRNA-incW-1 | tacttgctgtaactacttgc | Curing incW-oriV derived plasmids | This work |
| crRNA-incW-2 | ggatagtttttggagcgggc | Curing incW-oriV derived plasmids | This work |
| crRNA-incW-3 | gataactaccctctatccac | Curing incW-oriV derived plasmids | This work |

**Supporting Table S6. Primers.**

| Primer name | Sequence (5' to 3') | Purpose | Reference |
|---|---|---|---|
| VF2 | tgccacctgacgtctaagaa | Colony PCR, Sanger sequencing | 1, pSB1C3 |
| VR | attaccgcctttgagtgagc | Colony PCR, Sanger sequencing | 1, pSB1C3 |
| CmR-F | accgtaacacgccacatctt | qPCR primer of CmR | This work |
| CmR-R | ttcttgcccgcctgatgaat | qPCR primer of CmR | This work |
| AmpR-F | ttgatcgttgggaaccggag | qPCR primer of AmpR | This work |
| AmpR-R | tccgcctccatccagtctat | qPCR primer of AmpR | This work |
| KanR-F | tgatattcggcaagcaggca | qPCR primer of KanR | This work |
| KanR-R | gaccaccaagcgaaacatcg | qPCR primer of KanR | This work |
| SmR-F | ggctggctcgaagatacctg | qPCR primer of SmR | This work |
| SmR-R | ttggaaacttcggcttcccc | qPCR primer of SmR | This work |
| int 1-inversion-F | tagacgacttgaagacgtgacttgtggggc | Primer of int 1-based inversion PCR test. | This work |
| int 2-inversion-F | gacgagtcctgagagttccttggccttctt | Primer of int 2-based inversion PCR test. | This work |
| int 3-inversion-F | gggcttgagactacggatttttgcgtcgtt | Primer of int 3-based inversion PCR test. | This work |
| int 4-inversion-F | cacgaaaaacagggttttagttatacggctttagt tatacgtcagc | Primer of int 4-based inversion PCR test. | This work |

II. Supporting Results

A

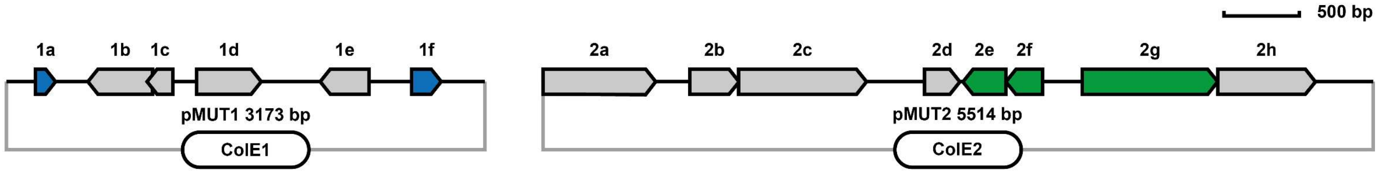

B

| ORFs | Range (bp) | Gene | Annotation | Notes |
|---|---|---|---|---|
| pMUT1 (GenBank: CP058218), 3173 bp | | | | |
| 1a | 194-325 | N/A | Replication initiation protein | Involved in plasmid replication |
| 1b | 541-969 | N/A | Hypothetical protein | Uncharacterized protein |
| 1c | 933-1103 | N/A | Hypothetical protein | Uncharacterized protein |
| 1d | 1261-1689 | N/A | Helix-turn-helix domain-containing protein | Uncharacterized protein |
| 1e | 2078-2401 | N/A | nikA protein | mobA, involved in plasmid mobilization |
| 1f | 2685-2879 | N/A | Rop family plasmid primer RNA-binding protein | Involved in plasmid copy number control |
| pMUT2 (GenBank: CP058219), 5514 bp | | | | |
| 2a | 3-749 | N/A | DUF4868 domain-containing protein | Uncharacterized protein |
| 2b | 976-1302 | N/A | MobC family plasmid mobilization relaxosome protein | mbeC, involved in plasmid mobilization |
| 2c | 1292-2140 | N/A | Relaxase/mobilization nuclease domain-containing protein | mbeA, involved in plasmid mobilization |
| 2d | 2526-2759 | mbeD | MbeD family mobilization/exclusion protein | mbeD, involved in plasmid mobilization |
| 2e | 2804-3088 | N/A | Type II toxin-antitoxin system RelE/ParE family toxin | toxin RelE |
| 2f | 3078-3320 | N/A | Type II toxin-antitoxin system RelB/DinJ family antitoxin | antitoxin RelB |
| 2g | 3585-4481 | N/A | Replication initiation protein | Replicase, involved in plasmid replication |
| 2h | 4471-5118 | N/A | Hypothetical protein | Uncharacterized protein |
| Gray background: redundant genes | | | | |

C

**pMUT2 oriV**
(26 bp, range: 751 bp - 776 bp)
**5' - acaaaacaacctatatcagataacag - 3'**

D

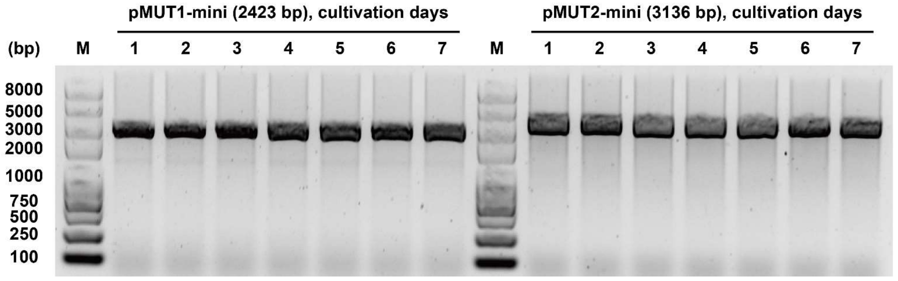

**Supporting Figure S1. Minimization and stability test of EcN cryptic plasmids pMUT1 and pMUT2.**

(**A**) Plasmid maps of pMUT1 and pMUT2.

(**B**) Details of pMUT1 and pMUT2.

(**C**) Identified pMUT2 (ColE2 group plasmid) origin of replication (oriV) sequence as 26 bp.

(**D**) Stability test of pMUT1-mini (pFB342) and pMUT2-mini (pFB344). Image of gel post electrophoresis showing the genetic stability of two minimized plasmids. Two *E. coli* Mach1-T1 strains harboring plasmids (pFB342 or pFB344) were continuously cultured (5 mL LB medium with antibiotics as 34 μg/mL chloramphenicol or 50 μg/mL kanamycin, 37 °C, 250 rpm, 16 h) in 7 days, and plasmids were extracted and digested by EcoRI each day. There was no size-variation in 7 days, showing the acceptable stability of two minimized plasmids.

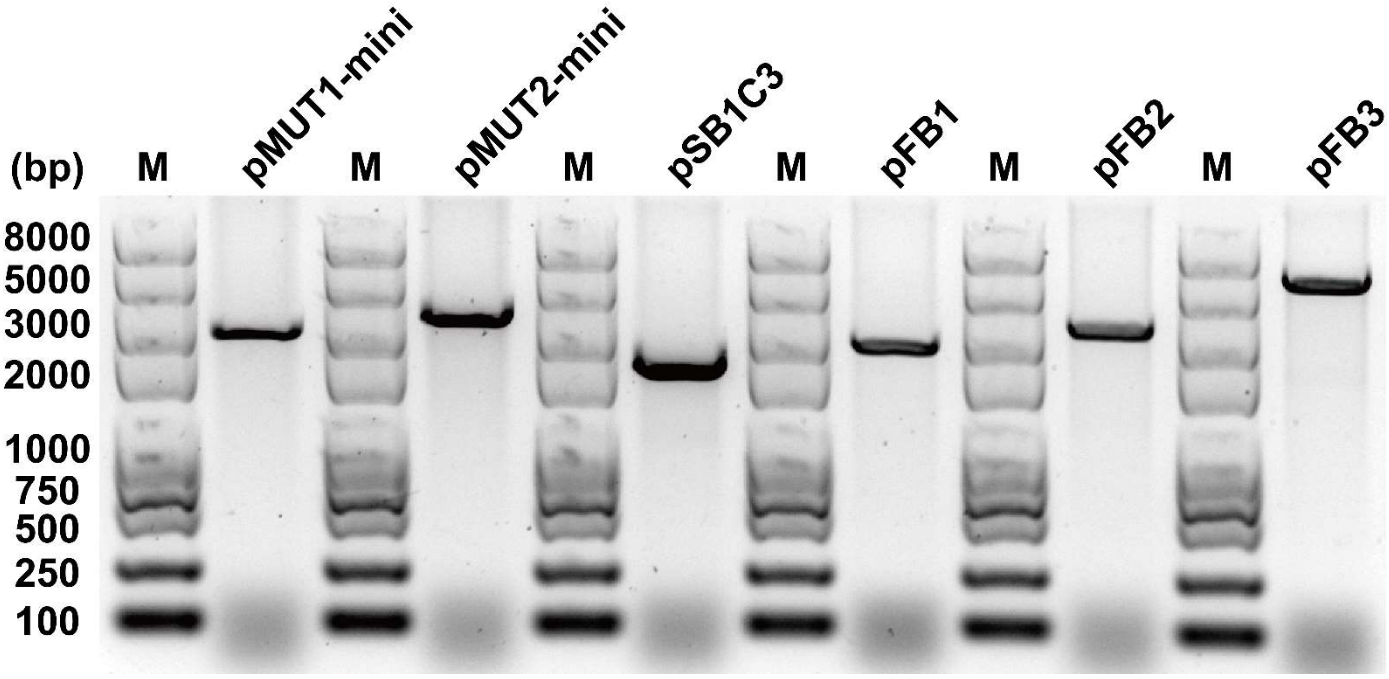

**Supporting Figure S2. Six compatible plasmid vectors.**

Agarose gel electrophoresis analysis of six compatible vectors that linearized by FastDigest XbaI: pMUT1-mini (pFB342, 2423 bp), pMUT2-mini (pFB344, 3136 bp), pSB1C3 (2070 bp), pFB1 (2333 bp), pFB2 (2771 bp), and pFB3 (5210 bp).

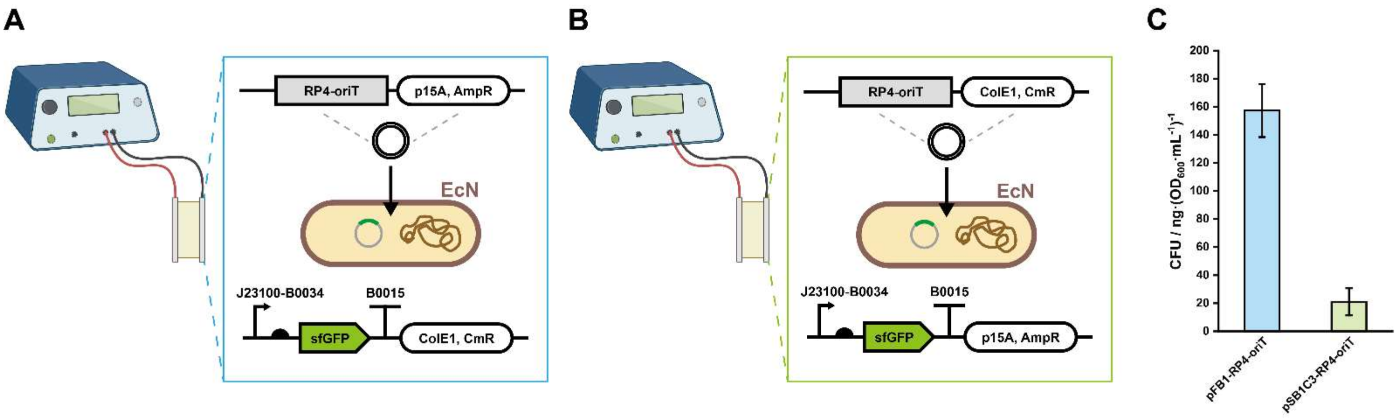

**Supporting Figure S3. Electroporation-based plasmid transformation of EcN.**

(**A**) pFB345 was transformed into EcN (harboring pFB346) via electroporation.

(**B**) pFB347 was transformed into EcN (harboring pFB348) via electroporation.

(**C**) Transformation efficiency of (**A**) and (**B**). Each value (mean ± standard deviation, SD) was calculated with three biological replicates.

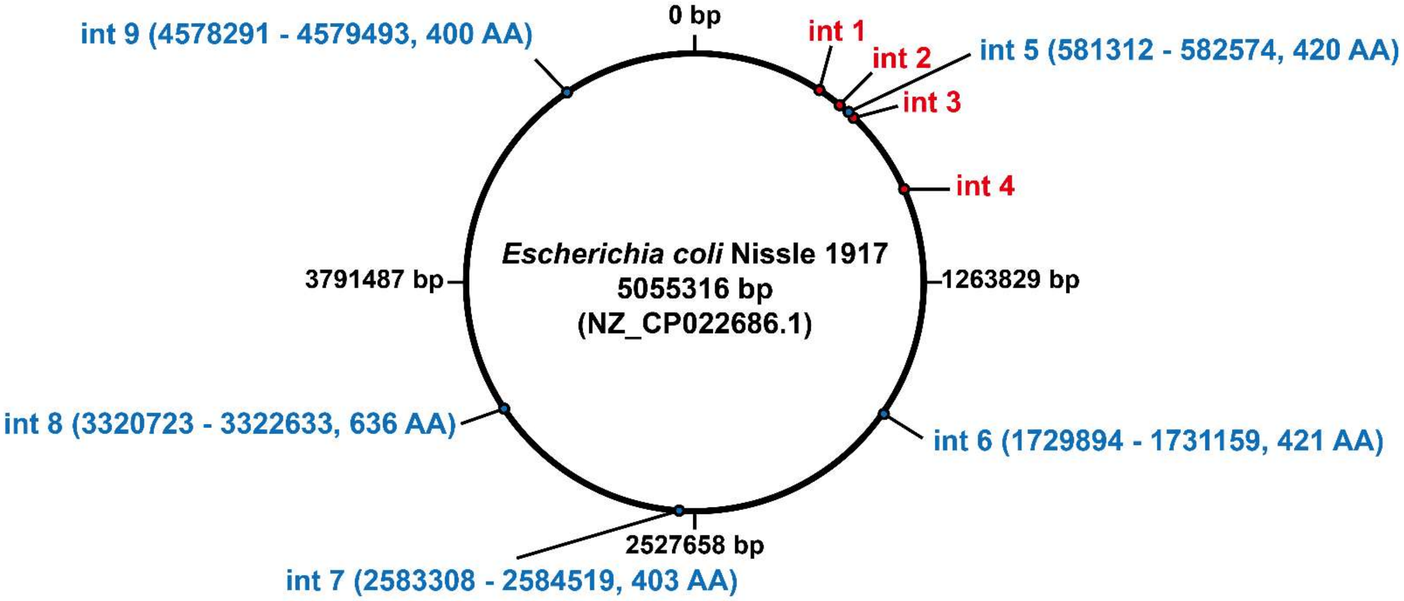

**Supporting Figure S4. Identification of EcN native integrases.**

int 1 to int 4 (red labelled) were successfully characterized in this study. The other five integrases (int 5 to int 9, blue labelled) were discovered, aligned and analyzed by InterPro, but their prophage sequence and *attP* / *attB* sites were not successfully identified in this study. (AA: amino acid)

| EcN-int 1 (437 AA) | | | | |
|---|---|---|---|---|
| | Accession | Name | Matches (AA) | Source database |
| Entries | IPR022000 | Min27-like integrase, DNA-binding domain | 7-70 | InterPro |
| | IPR010998 | Integrase/recombinase, N-terminal | 77-198 | InterPro |
| | IPR044068 | Core-binding (CB) domain | 82-176 | InterPro |
| | IPR011010 | DNA breaking-rejoining enzyme, catalytic core | 82-401 | InterPro |
| | IPR002104 | Integrase, catalytic domain | 197-409 | InterPro |
| | IPR013762 | Integrase-like, catalytic domain superfamily | 201-413 | InterPro |
| Residues | cd01189 | C-terminal catalytic domain of integrases from bacterial phages and conjugate transposons | 232R, 359Y, 362R, 363H, 395Y | CDD |

| EcN-int 2 (406 AA) | | | | |
|---|---|---|---|---|
| | Accession | Name | Matches (AA) | Source database |
| Entries | IPR038488 | Integrase, DNA-binding domain superfamily | 6-115 | InterPro |
| | IPR025166 | Integrase, DNA-binding domain | 22-101 | InterPro |
| | IPR044068 | Core-binding (CB) domain | 113-194 | InterPro |
| | IPR011010 | DNA breaking-rejoining enzyme, catalytic core | 114-397 | InterPro |
| | IPR010998 | Integrase/recombinase, N-terminal | 116-196 | InterPro |
| | IPR013762 | Integrase-like, catalytic domain superfamily | 223-393 | InterPro |
| | IPR002104 | Integrase, catalytic domain | 225-397 | InterPro |
| Residues | cd00801 | Bacteriophage P4 integrase, C-terminal catalytic domain | 262R, 347H, 350R, 351S, 384Y | CDD |

| EcN-int 3 (423 AA) | | | | |
|---|---|---|---|---|
| | Accession | Name | Matches (AA) | Source database |
| Entries | IPR038488 | Integrase, DNA-binding domain superfamily | 1-87 | InterPro |
| | IPR025166 | Integrase, DNA-binding domain | 3-87 | InterPro |
| | IPR010998 | Integrase/recombinase, N-terminal | 89-200 | InterPro |
| | IPR044068 | Core-binding (CB) domain | 96-176 | InterPro |
| | IPR011010 | DNA breaking-rejoining enzyme, catalytic core | 98-389 | InterPro |
| | IPR002104 | Integrase, catalytic domain | 199-392 | InterPro |
| | IPR013762 | Integrase-like, catalytic domain superfamily | 201-397 | InterPro |
| Residues | cd00801 | Bacteriophage P4 integrase, C-terminal catalytic domain | 239R, 342H, 345R, 346A, 379Y | CDD |

| EcN-int 4 (413 AA) | | | | |
|---|---|---|---|---|
| | Accession | Name | Matches (AA) | Source database |
| Entries | IPR038488 | Integrase, DNA-binding domain superfamily | 7-94 | InterPro |
| | IPR025166 | Integrase, DNA-binding domain | 8-94 | InterPro |
| | IPR010998 | Integrase/recombinase, N-terminal | 96-210 | InterPro |
| | IPR044068 | Core-binding (CB) domain | 105-187 | InterPro |
| | IPR011010 | DNA breaking-rejoining enzyme, catalytic core | 105-386 | InterPro |
| | IPR002104 | Integrase, catalytic domain | 210-387 | InterPro |
| | IPR013762 | Integrase-like, catalytic domain superfamily | 211-388 | InterPro |
| Residues | cd00801 | Bacteriophage P4 integrase, C-terminal catalytic domain | 249R, 338H, 341R, 342A, 374Y | CDD |

**Supporting Figure S5. InterPro annotation of four EcN native integrases.**

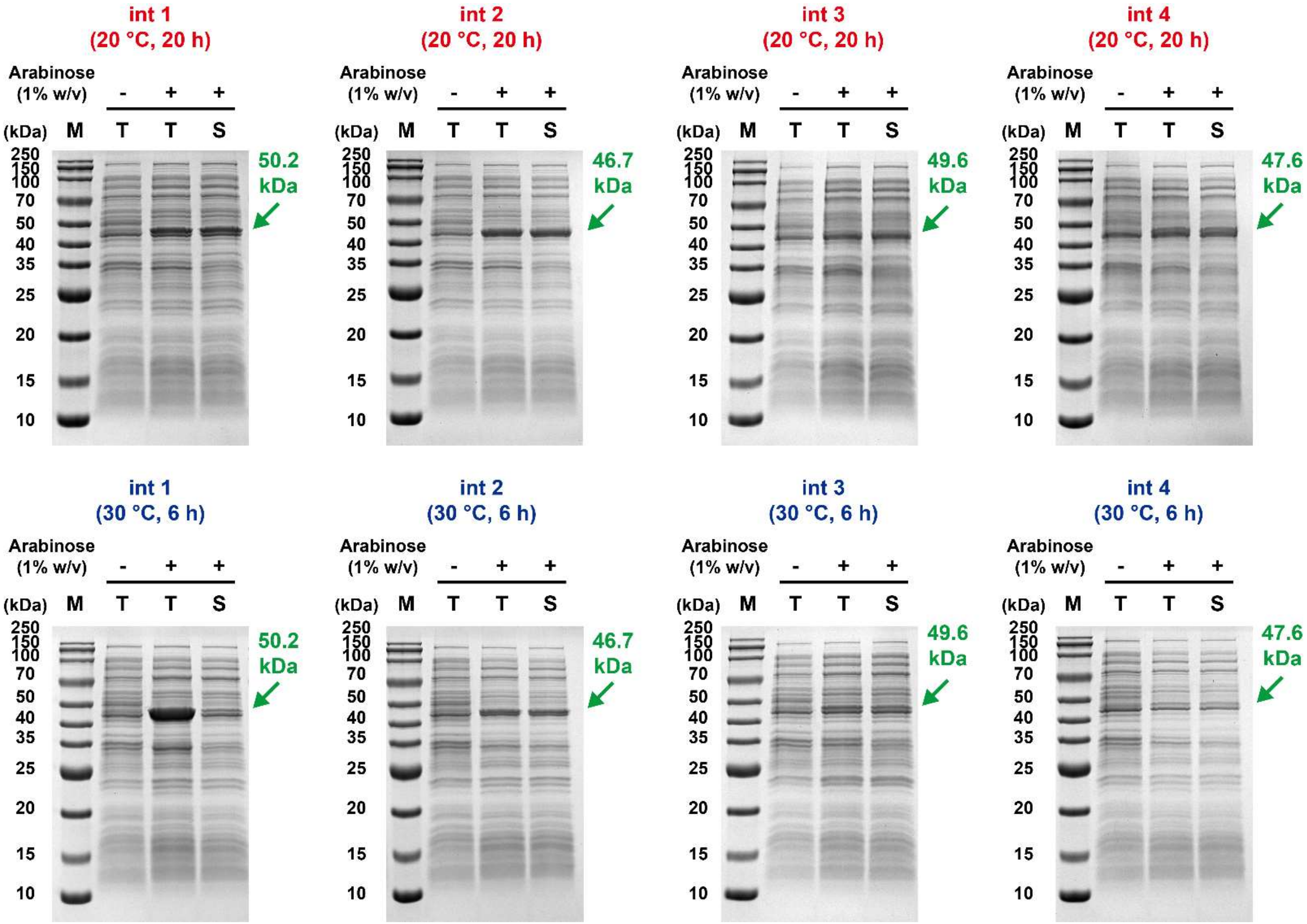

**Supporting Figure S6. *In vivo* expression of four EcN native integrases.**

The *in vivo* protein expression was performed as the same as our previous work.[2] The difference is that pFB360-pFB363 were used in *E. coli* Mach1-T1 for integrase expression, and the inducer was replaced by 1% w/v arabinose than 0.5 mM IPTG. The results showed that a lower (20 °C for 20 h) temperature was preferred for soluble integrase expression.

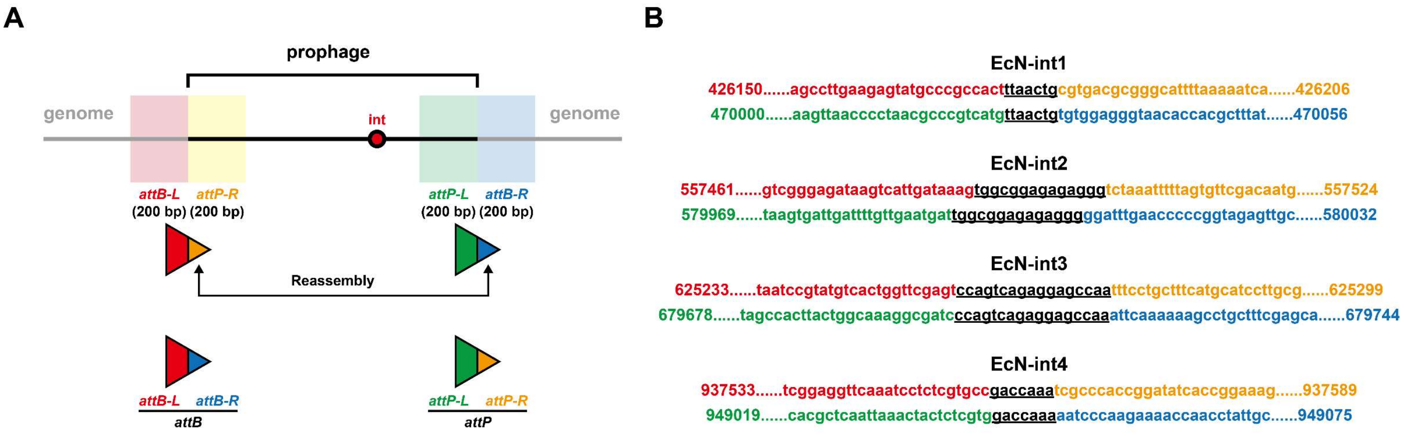

**Supporting Figure S7. Reassembly of *attP* / *attB* sites.**

(**A**) Workflow of *attP* / *attB* reassembly of four EcN native integrases.

(**B**) DNA sequence and location of *att* sites in EcN genome. The reassembled sites were constructed into pFB364-pFB367, and successfully demonstrated their function. The core sequences were labelled as **underlined black** (note that the two core sequences for one integrase are the same). The other sequences (25 bp for each, partially) were shown as *attB-L*, *attP-R*, *attP-L*, and *attB-R*, respectively.

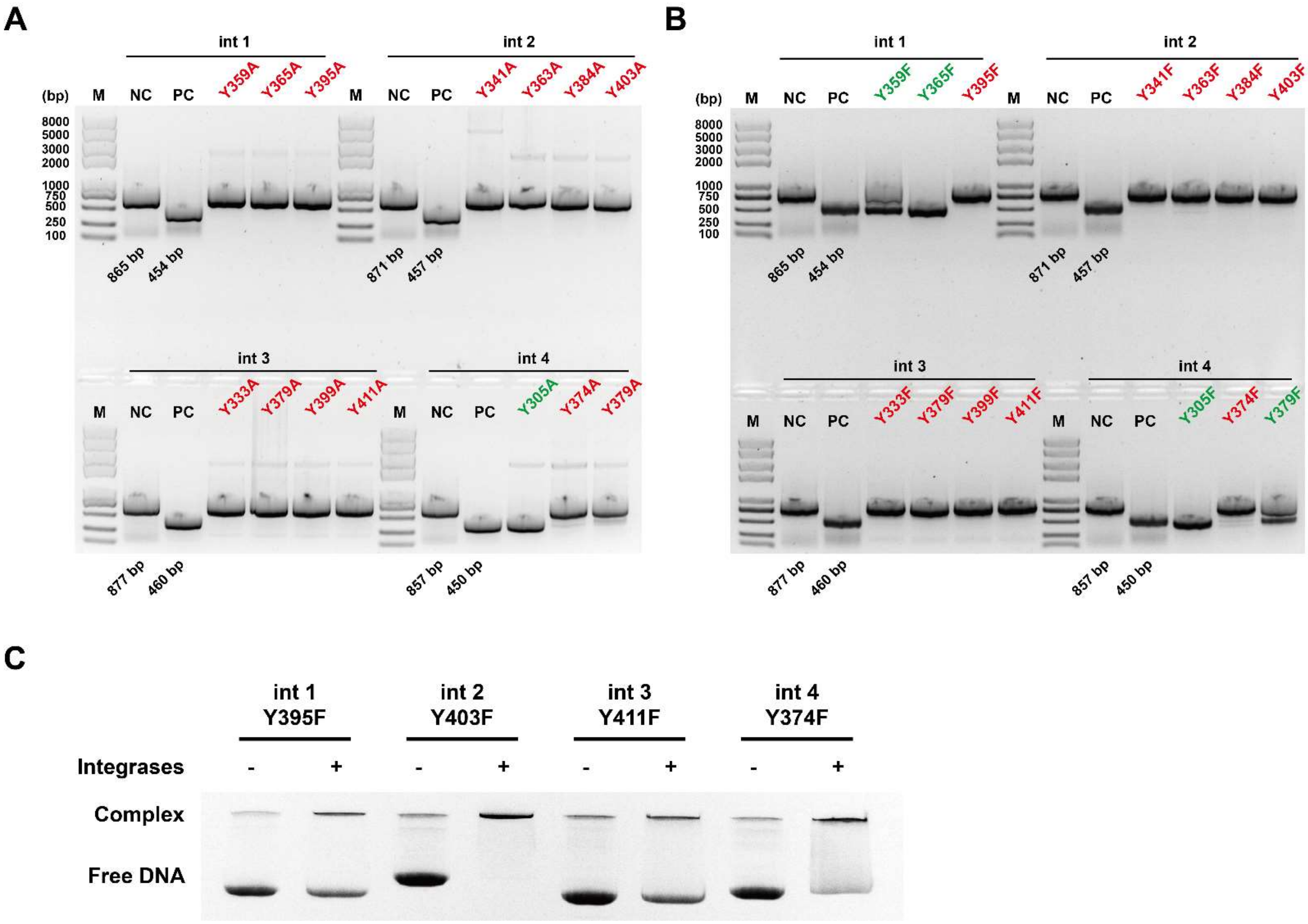

**Supporting Figure S8. Tyrosine residue mutation of four EcN native integrases.**

(**A**) C-terminal tyrosine (Y) residues were mutated into alanine (A) and then test their activity. Red-labelled variants indicate the inactivation, and green-labelled variants indicate the activation still remains. The PCR products were amplified from plasmids pFB364-pFB367 (contain *attP* / *attB* sites).

(B) C-terminal tyrosine (Y) residues were mutated into phenylalanine (F) as previous reported,[5] and then test their activity. Red-labelled variants indicate the inactivation, and green-labelled variants indicate the activation still remains. The PCR products were amplified from plasmids pFB364-pFB367 (contain *attP* / *attB* sites).

(**C**) Electrophoretic Mobility Shift Assay (EMSA) results showed that inactivated integrases still remain their DNA-binding capability.

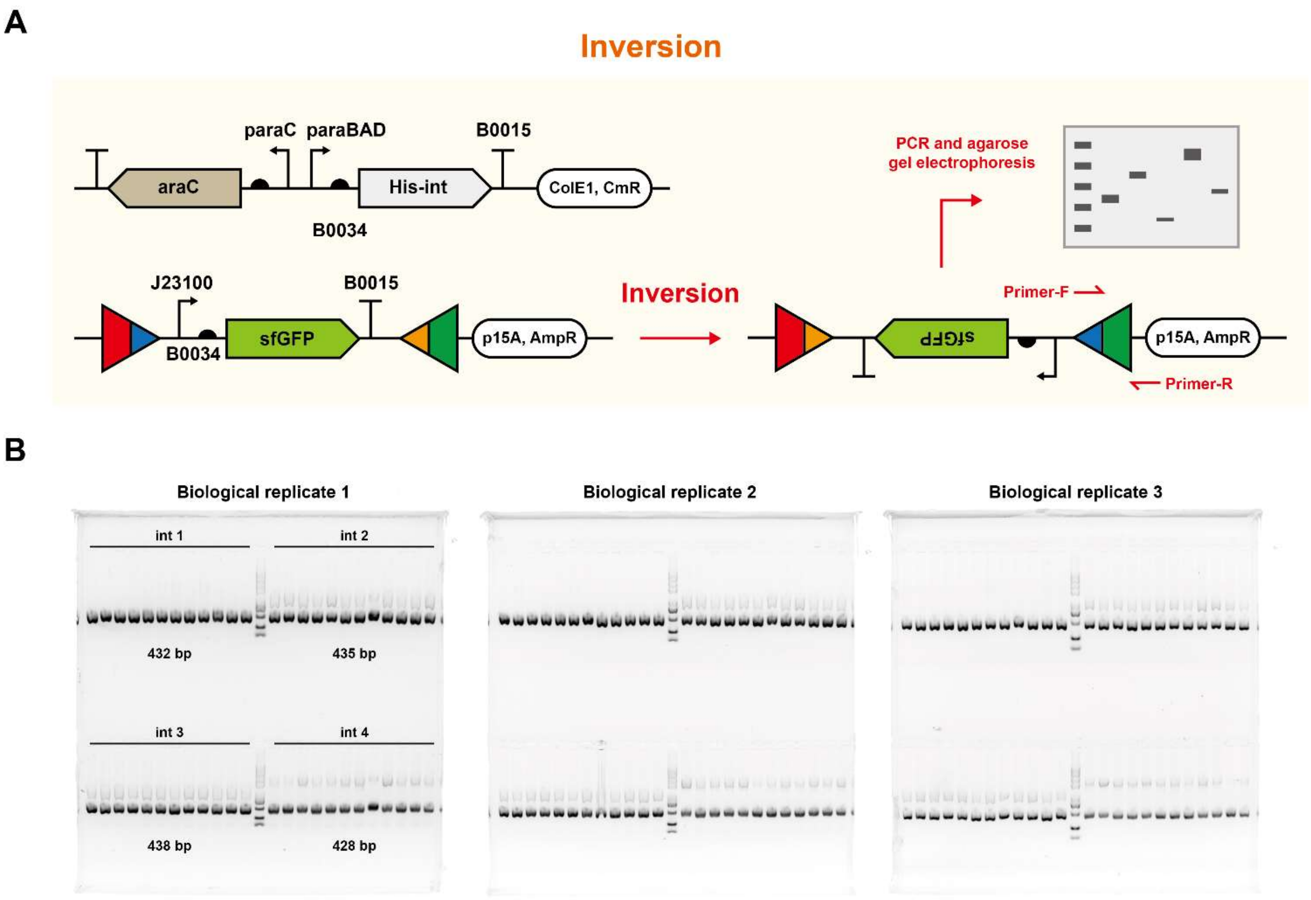

**Supporting Figure S9. Inversion test of four EcN native integrases.**

(**A**) Schematic diagram of inversion test for four EcN native integrases.

(**B**) The inversion efficiency was calculated by PCR-agarose gel electrophoresis (12 colonies were picked up and test for one integrase each time) with three biological replicates.

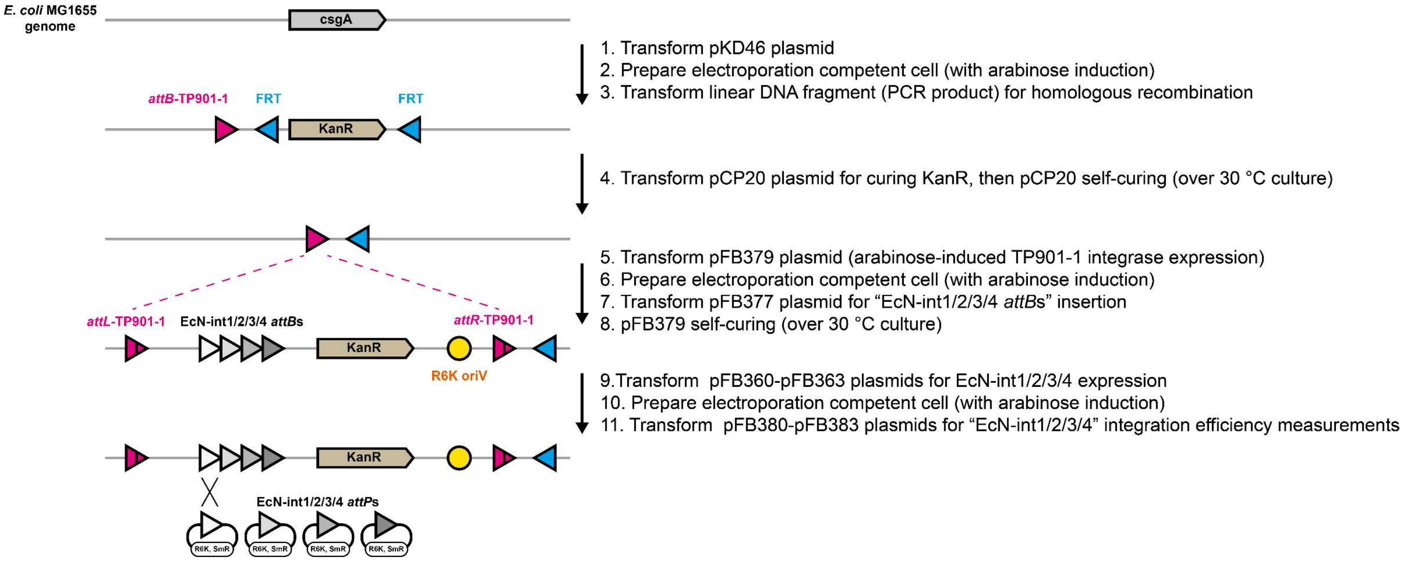

**Supporting Figure S10. Workflow of *attBs* (four EcN native integrases) integration into *E. coli* MG1655 genome.**

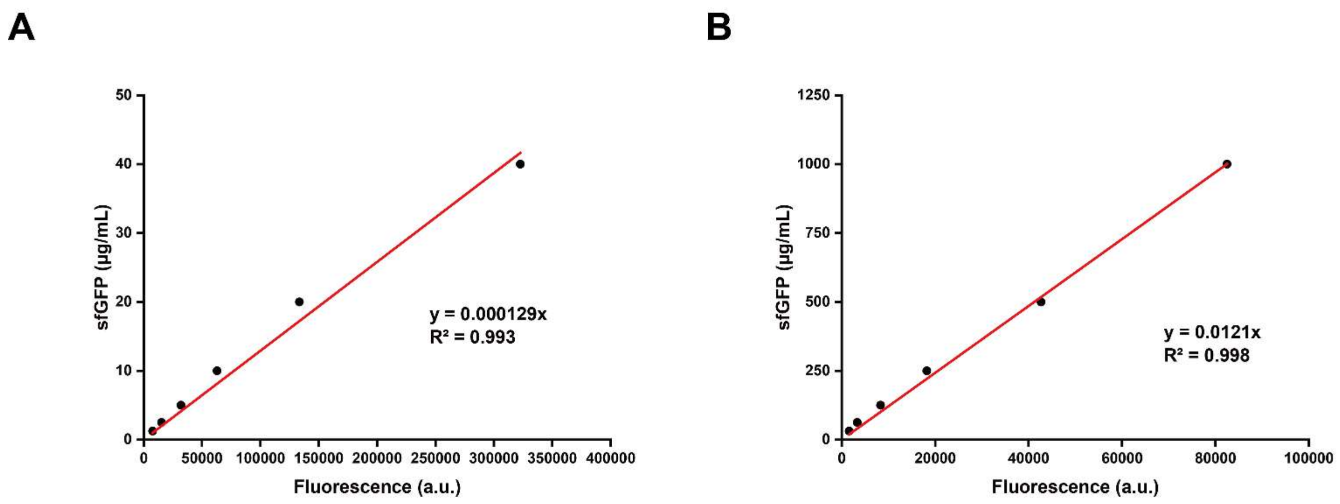

**Supporting Figure S11. Standard curves of "sfGFP yield-Fluorescence" conversion.**

(**A**) Conversion curve for *in vivo* sfGFP conversion.

(**B**) Conversion curve for *in vitro* CFPS sfGFP conversion.

III. Supporting Methods

**Preparation of EcN CFPS crude extract**

**Day 1:**

- Note: Here the EcN extract is prepared in 1 L batches.

1. Prepare 1 L sterile 2×YTPG liquid medium in a 2.5 L flask.

    - For 1 L 2×YTPG liquid medium preparation, add:

    a. (in 800 mL water for sterilization) 10 g yeast extract, 16 g tryptone, 5 g sodium chloride (NaCl), 7 g dipotassium phosphate ($K_2HPO_4$), 3 g monopotassium phosphate ($KH_2PO_4$).

    b. (in 200 mL water for sterilization) 18 g glucose.

    Adjust “a” to pH 7.2 before sterilization (by potassium hydroxide, KOH), and mix “a” with “b” after sterilization, if the volume does not reach to 1 L, then add sterile water to 1 L for the following cultivation.

2. Incubate 25 mL LB liquid medium (in 250 mL flask) overnight (with EcN strain, which is picked up from LB-agar plate colonies or transferred from LB liquid medium) with antibiotics at 37 °C, 250 rpm for about 16 h.

**Day 2:**

1. Add the 25 mL EcN culture to 1 L 2×YTPG medium, incubate at 37 °C, 250 rpm for about 3 h, the goal of $OD_{600}$ is about 3.

    - To ensure that $OD_{600}$ does not exceed 3, measure the value every 30 min.

2. While waiting for $OD_{600}$ reaches to 3, prepare for the centrifugation steps.

    - Prepare 125 mL S30 buffer (125 mL / batch) in a flask, bury in ice when done, the 125 mL S30 buffer contains: 1.25 mL 1 M Tris-acetate, 1.25 mL 6 M potassium acetate, 1.25 mL 1.4 M magnesium acetate, 0.25 mL 1 M dithiothreitol (DTT).

3. When $OD_{600}$ = 3, collect the EcN pellets by centrifugation at 5000 g, 4 °C for 15 min.

4. Resuspend the pellets as “10 mL S30 buffer + 1 g wet weight EcN pellet”, and collect the mixtures into several 50 mL centrifuge tubes.

5. centrifuge tubes at 10000 g, 4 °C for 15 min.

    - After once and more resuspension, the pellets will become loose and hard to sediment, so here we increase the centrifugal force from 5000 g to 10000 g.

6. Discard the supernatant, resuspend the pellets as “10 mL S30 buffer + 1 g wet weight EcN pellet”, and centrifuge tubes at 10000 g, 4 °C for 15 min.

7. repeat step “6”.

8. Discard the supernatant, dry tubes completely, reweight the pellets and label the mass on the tubes, then flash freeze in liquid nitrogen, store in -80 °C freezer for later uses.

**Day 3:**

1. Remove tubes from -80 °C freezer to ice and thaw for about 60 min.

2. Resuspend pellets as “1 mL S30 buffer + 1 g wet weight EcN pellet”, and transfer 1.4 mL mixtures into new 1.5 mL centrifuge tubes.

3. Sonicate mixtures in ice-water mixture as the parameters: 10 seconds on / 10 seconds off, 50% amplitude, 1000 Joule / mL. Once finished the sonication, add 4 μL 1 M DTT and invert gently to mix.

4. Centrifuge the lysis at 20000 g, 4 °C for 15 min.

5. After the centrifugation, the mixture will be shown as three layers, remove the top layer (about 600 µL supernatant) from each tube and transfer the top layer individually into a new 1.5 mL centrifuge tube.

- For EcN cell extract preparation, the run-off step is not necessary. Our experiment showed that after a run-off reaction (the supernatant from “5” were incubated at 37 °C for another 60 min), the extract will totally lose activity (data not shown).

6. Centrifuge the supernatant again at 20000 g, 4 °C for 15 min, aiming to discard the residual pellets.

7. After centrifugation, remove 500 µL supernatant and transfer it into a new 1.5 mL tube as the final EcN cell extract. Label the tubes correctly and stored in -80 °C freezer for later uses.
